# Introns encode a vast new class of Kink-loop RNAs that autoregulate pre-mRNA splicing

**DOI:** 10.64898/2026.08.03.742641

**Authors:** Bin Li, Qiao Lin, Anrui Liu, Haofei Gan, Shurong Liu, Wujian Zheng, Yonglin Liang, Junhong Huang, Keren Zhou, Gang Wan, Lianghu Qu, Jianhua Yang

**Affiliations:** MOE Key Laboratory of Gene Function and Regulation, State Key Laboratory of Biocontrol, Guangdong Provincial Key Laboratory of Pharmaceutical Functional Genes, Center of Evolutionary Synthetic Biology, Sun Yat-sen University, Guangzhou, Guangdong, China; Hong Kong Institute of Advanced Studies, Sun Yat-sen University, Hong Kong, China; Koch Institute for Integrative Cancer Research, Massachusetts Institute of Technology, 500 Main Street, Cambridge, MA 02139, USA; Department of Pathology, St. Jude Children’s Research Hospital, 262 Danny Thomas Place, Memphis, TN, 38105, USA

**Keywords:** RNA structure motif, Kink-loop, klRNA, 15.5K, intron splicing, intron autoregulation

## Abstract

Introns occupy nearly one-third of the human genome, yet whether they encode widespread regulatory functions remains unclear. Here, we identify a vast new class of intron-derived Kink-loop RNAs (klRNAs) that autoregulate host pre-mRNA splicing. We developed orthogonal sequencing methods to systematically uncover approximately 15,200 and 3,500 previously unannotated klRNAs in humans and mice, respectively. Bound by the conserved RNA-binding protein 15.5K, klRNAs are compact orphan RNAs characterized by stereotypically positioned terminal C/D motifs that form K-loop structures. Depletion of 15.5K broadly disrupts klRNA biogenesis. Functional and genetic perturbations establish that klRNAs suppress host intron excision through a K-loop-dependent mechanism. Together, our findings establish klRNA-mediated autoregulation as a widespread principle governing intron fate and reveal a previously unrecognized regulatory layer encoded within mammalian introns.

## Main

The evolutionary expansion of intronic sequence is a defining feature of complex eukaryotic genomes, yet the functional potential of this vast non-coding sequence space remains unclear ^1, 2^. In humans, introns occupy nearly one-third of the genome and constitute a major source of transcribed non-coding sequences ^1, 2^. Although introns give rise to diverse functional non-coding RNAs (ncRNAs), including microRNAs (miRNAs), small nucleolar RNAs (snoRNAs), and long non-coding RNAs (lncRNAs), these annotated RNA classes account for only a small fraction of intronic sequence space ^3–5^. Whether introns encode previously unrecognized regulatory RNA systems and how such systems contribute to gene expression remain fundamental questions in genome biology.

Structured RNA motifs provide a versatile means of encoding biological information through defined RNA architectures and protein interactions. Among these structure motifs, Kink-turn (K-turn) motifs represent evolutionarily conserved RNA tertiary architectures that recruit members of the L7Ae/15.5K protein family and promote ribonucleoprotein assembly across diverse organisms ^6–8^. Canonical K-turns consist of two RNA stems connected by a short asymmetric internal loop that introduces a characteristic kink in the RNA backbone ^6–8^. A related but structurally distinct architecture, the Kink-loop (K-loop), was subsequently identified in archaeal box C/D RNAs (cdRNAs), where loss of the canonical C-stem allows internal C′/D′ motifs to form K-loop structures that retain recognition by L7Ae proteins ^9^. However, known K-loops have remained largely confined to internal C′/D′ motifs within canonical cdRNA scaffolds. Whether K-loop architecture can be deployed beyond this canonical context to define distinct RNA classes with alternative regulatory functions remains unknown.

In this study, we developed two orthogonal sequencing methods, enhanced RIP-PEN-seq (eRIP-PEN-seq) and D-box-enriched full-length RNA sequencing (CD-seq), to systematically identify 15.5K-associated structured ncRNAs, uncovering a vast new class of Kink-loop RNAs (klRNAs). We identify approximately 15,200 and 3,500 novel klRNAs in humans and mice, respectively, revealing one of the largest classes of structured ncRNAs encoded in mammalian genomes. Unlike canonical cdRNAs, klRNAs are compact, predominantly intron-derived orphan RNAs that generally lack canonical antisense targets and form K-loop structures through terminal C/D motifs rather than internal C′/D′ motifs of canonical archaeal cdRNAs. Mechanistically, klRNAs require 15.5K for their biogenesis and act predominantly *in cis* to suppress host intron excision through a K-loop-dependent mechanism, establishing klRNA-mediated intron autoregulation as a widespread regulatory principle.

## Results

### Introns encode a vast new class of K-loop RNAs bound by 15.5K

To systematically identify endogenous and full-length RNAs bound by 15.5K protein, we developed an enhanced RIP-PEN-seq (eRIP-PEN-seq, **Extended Data Fig. 1a, details in Methods**), which depletes highly abundant ncRNAs (rRNAs, snRNAs and snoRNAs) to improve recovery of low-abundance transcripts. A total of approximately 162 million paired-end reads were generated from eRIP-PEN-seq **(Supplementary Table 1)**. Compared to our previous RIP-PEN-seq protocol ^10^, where 1.3% of reads aligned to unannotated genomic regions (including intronic and intergenic regions), eRIP-PEN-seq substantially improved specificity, with 11.0% of reads mapping to unannotated regions **(Extended Data Fig. 1b, and Supplementary Table 1)**. This enhanced enrichment of novel RNAs suggests that eRIP-PEN-seq effectively reduces interference from abundant ncRNAs, thereby improving the detection of previously uncharacterized transcripts.

We next developed a novel pipeline to identify transcript units (TUs) by clustering overlapping aligned reads (**Extended Data Fig. 1c**). We identified 14862 (78%) novel TUs, the majority of which originated from intronic regions (66.3%), along with a smaller fraction from intergenic loci (2227, 11.7%), as well as 4204 TUs that overlapped with known annotations, including 523 snoRNAs and 3681 other known annotations (**Extended Data Fig. 1d**). *De novo* motif analysis of these novel TUs revealed the significant enrichment of canonical C box motif (UGAUGA, P value = 1e-1804) and D box motif (CUGA, P value = 1e-744) **(Extended Data Fig. 1e)**. Moreover, the C box motif is primarily located near the 5’ end of RNA **(Extended Data Fig. 1f)**, while the D box motif is primarily located near the 3’ end **(Extended Data Fig. 1g)**. These observations indicate that our eRIP-PEN-seq data may harbor a large number of yet-unidentified intron-encoded cdRNAs.

We further applied our computational method cdSeeker ^11^ (**Extended Data Fig. 1h**) to identify novel cdRNAs from the TUs of 15.5K eRIP-PEN-seq data by scoring the C box and D box motifs. We retained novel cdRNAs with a score of 11.5 bits or higher and further filtered candidates requiring presence in at least two sequencing libraries. This analysis identified 9,135 high-confidence novel cdRNAs in the 15.5K eRIP-PEN-seq datasets **(Extended Data Fig. 1i and Supplementary Table 2)**, more than 85% of which originated from intronic regions **(Fig. 1a)**. By comparison, application of the same analysis framework to the previous 15.5K RIP-PEN-seq datasets identified 5,680 cdRNAs **(Extended Data Fig. 1j and Supplementary Table 2)**. Thus, eRIP-PEN-seq substantially increased the recovery of 15.5K-associated cdRNAs relative to RIP-PEN-seq, demonstrating its enhanced sensitivity for detecting this previously underrepresented RNA population.

**Figure 1.**
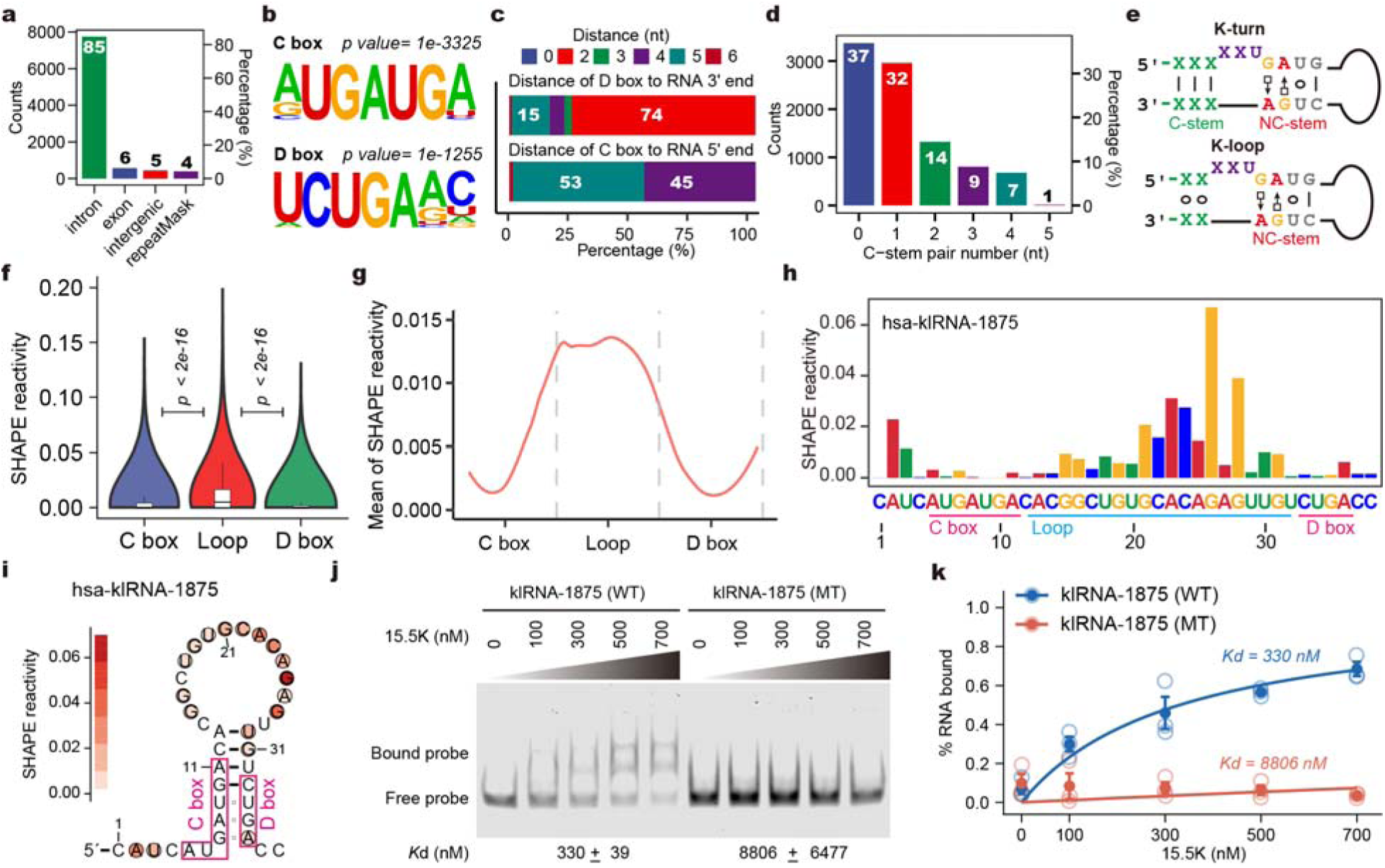
Discovery of a vast class of intron-derived K-loop RNAs recognized by 15.5K. **a,** Genomic distribution of newly identified cdRNAs recovered by eRIP-PEN-seq. **b,** Enriched motifs identified in human novel cdRNAs by HOMER software. Two significant motifs (C box: RUGAUGA; D box: CUGA) were identified in the RNAs. **c,** Positioning of C-box and D-box motifs relative to RNA termini. **d,** Distribution of C-stem base-pair numbers among newly identified cdRNAs. **e,** Comparison of canonical K-turn RNAs (ktRNAs) and newly identified K-loop RNAs (klRNAs). Unlike canonical K-turn RNAs (ktRNAs), klRNAs lack the canonical C-stem while retaining the NC-stem and loop architecture characteristic of K-loop formation. X represents any nucleotide. **f,** SHAPE reactivity across klRNAs, including the C box, loop region, and D box, averaged across all klRNAs. The box plots indicate the median and the upper and lower quartiles. The P values of the differences in distance between the two categories were determined by the Mann-Whitney-Wilcoxon test. **g,** Metagene profile of SHAPE reactivity across klRNAs. **h,** The SHAPE reactivity signal on hsa-klRNA-1875. The C and D boxes and the loop region are marked with pink and blue underlines in the bar plot, respectively. The SHAPE reactivity is calculated from merged n= 4 biological replicates. **i,** The predicted secondary structure of the K-loop on hsa-klRNA-1875. The C and D boxes are indicated with pink boxes in the structure figures. The SHAPE reactivity is calculated from merged n= 4 biological replicates. **j, k,** RNA EMSA (**j**) showing specific binding of recombinant 15.5K protein to wild-type (WT) but not K-loop mutant (MT) klRNA. Quantification of independent EMSA (**k**) reveals the percentage of bound probes with increasing concentration of 15.5K over three replicates for WT- and MT-klRNA-1875, respectively. The binding curves are presented as mean ± s.d. (n = 3 for each data point). The dissociation constant (Kd, nM) values were indicated at the lower panel of the REMSA image.

Intriguingly, *de novo* motif enrichment revealed a strikingly stereotyped terminal organization, with a C-box motif (RUGAUGA) positioned predominantly 4 (45%) or 5 nucleotides (53%) downstream of the 5′ terminus and a D-box motif (CUGA) located predominantly 2 (74%) or 5 nucleotides (15%) upstream of the 3′ terminus (**Fig. 1b, c and Extended Data Fig. 1k**). Given that C- and D-box motifs at RNA termini are known to form K-turn structures, typically comprising a canonical C-stem with at least two base pairs ^12^ and a non-canonical NC-stem, we investigated terminal base-pairing in these newly identified cdRNAs. Notably, only 31% of cdRNAs exhibited C-stems with two or more base pairs, consistent with canonical K-turn RNAs (ktRNAs) ^10^ (**Fig. 1d, e**). In contrast, 69% of cdRNAs contained fewer than two terminal base pairs (**Fig. 1d**), indicating that most of these RNAs lack the canonical C-stem required for K-turn architecture. This raised the possibility that their terminal C/D motifs instead form K-loop structures, analogous to the C-stem-lacking K-loops formed by internal C′/D′ motifs in canonical cdRNAs ^9^. We therefore define this expanded RNA class as K-loop RNAs (klRNAs), distinguished by K-loop architectures formed by precisely positioned terminal C/D motifs rather than the internal C′/D′ motifs of canonical archaeal cdRNAs. (**Fig. 1e, and Supplementary Table 2**).

To determine whether these C-stem-lacking klRNAs indeed adopt K-loop structures *in vivo*, we probed their intact RNA conformations using 15.5K RIP-PEN-SHAPE-MaP data. Paired nucleotides forming the NC-stem within the C and D boxes exhibited low SHAPE reactivity, whereas unpaired nucleotides in the loop region of novel klRNAs showed relatively high reactivity **(Fig. 1f and Supplementary Table 3).** Consistently, metagene analysis across klRNAs revealed reduced SHAPE reactivity within the C and D boxes, supporting their structured configuration **(Fig. 1g)**. For instance, the SHAPE reactivity profile of hsa-klRNA-1875 closely corroborated its predicted K-loop structure (**Fig. 1h and 1i**). Together, these structural data support the formation of K-loop conformations by klRNAs *in vivo*.

To further elucidate the RNA-binding properties of 15.5K toward klRNAs, we employed RNA electrophoretic mobility shift assays (REMSA) *in vitro*. Consistent with the binding observed for the canonical K-turn RNA hsa-ktRNA-930 **(Extended Data Fig. 1l-o)**, recombinant 15.5K selectively bound wild-type hsa-klRNA-1875, whereas disruption of its K-loop by a CUGA-to-CUAG mutation markedly reduced binding **(Fig. 1j and 1k)**. Together with the SHAPE-MaP data, these results support the formation of terminal C/D motif–defined K-loop structures in klRNAs and their direct recognition by 15.5K.

Applying cdSeeker to 15.5K RIP-PEN-seq data from mouse Hepa1-6 cells identified an additional 3,053 klRNAs **(Extended Data Fig. 2a and Supplementary Table 4)**. These mouse klRNAs exhibited genomic distributions and motif architectures similar to those observed in human klRNAs **(Extended Data Fig. 2b-f)**, indicating that this RNA class is broadly represented across mammals.

Together, these results identify a vast and previously unrecognized class of intron-derived K-loop RNAs (klRNAs) in mammalian genomes, characterized by terminal C/D motif–formed K-loop architectures and direct association with 15.5K.

### Introns encode thousands of novel klRNAs uncovered by CD-seq

To overcome the limitations of antibody-dependent enrichment in eRIP-PEN-seq, which relies on high-quality antibodies and large cell numbers, we developed D-box-enriched full-length RNA sequencing (CD-seq), a motif-anchored approach that selectively captures RNAs bearing terminal box D motifs **(Fig. 2a)**. This approach enables efficient enrichment of full-length cdRNAs through motif-anchored ligation and directional amplification, and was further adapted into CD2-seq and CD5-seq to accommodate the variable spacing between D-box termini and downstream nucleotides (see Methods, **Fig. 2b, and Extended Data Fig. 3a, b)**.

**Figure 2.**
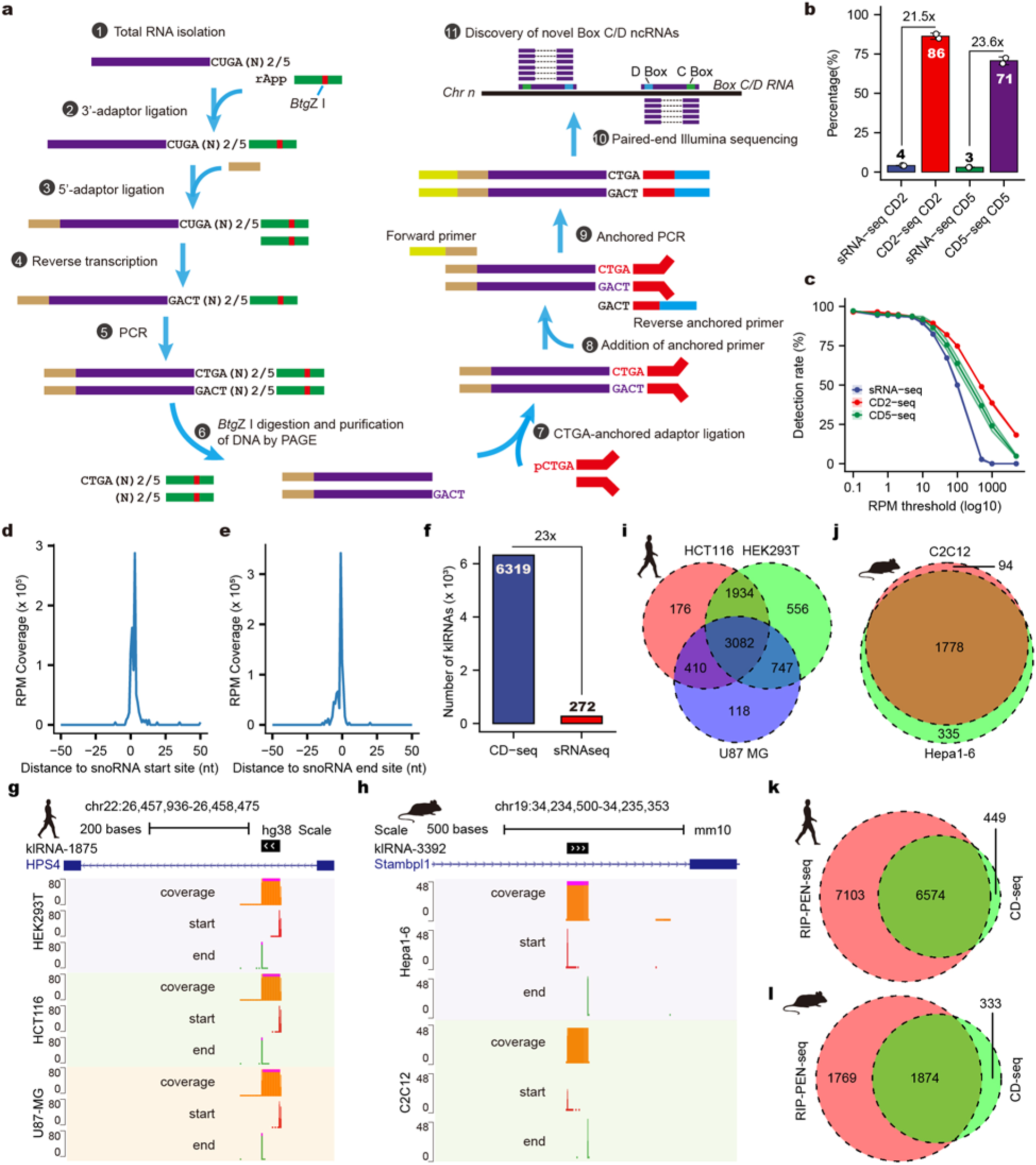
CD-seq enables transcriptome-wide discovery of thousands of novel klRNAs. **a,** Overview of the CD-seq strategy for selective enrichment of full-length D-box-containing RNAs. (1) Total RNA is extracted and treated with DNase I to remove genomic DNA. (2) Then a preadenylated 3’ adaptor containing a *Btg*Z I site (black line) is ligated to the 3’ end of the RNA. (3) After digestion of the remaining 3’ adaptor, the 5’ adaptor is attached to capture the information at the 5’ end of the RNA. (4) After the adaptor-ligation step, the ligated RNA is reverse transcribed using a specific primer complementary to the 3’ adaptor. (5) The cDNA is amplified by low-cycle PCR. (6) Subsequently, the double-stranded DNA is cleaved with *Btg*Z I and separated via native PAGE. The DNA in the range of 50 bp to 700 bp is recovered. (7) The motif-anchored adaptor is annealed with two oligos and ligated to the purified DNA with T4 DNA ligase. (8-9) The ligation products with standard Illumina sequences on both ends are amplified with the forward primer and reverse motif-anchored primer, which are both compatible with the Illumina system. (10) Paired-end sequencing is performed with the standard Illumina primer. (11) CD-seq reads are aligned to genomic sequences for further discovery of novel RNAs. **b,** Enrichment efficiency of D-box-containing RNAs compared with sRNA-seq. Values represent the mean ± s.e.m (n=2). **c,** Detection sensitivity of annotated box C/D RNAs by CD-seq and sRNA-seq. **d, e,** Representative loci illustrating accurate start (**d**) and end (**e**) mapping of canonical box C/D RNAs by CD-seq. **f,** Numbers of klRNAs identified by CD-seq and sRNA-seq. **g, h,** The UCSC genome browser displaying the CD-seq signals (coverage, orange; 5’-start, red; 3’-end, green) of two klRNAs in humans (**g**) and mice (**h**). **i,** Overlap analysis of klRNAs identified from human HEK293T, HCT116 and U87-MG cells. **j,** Overlap analysis of klRNAs identified from mouse C2C12 and Hepa1-6 cells. **k, l,** Comparison of klRNAs identified from RIP-PEN-seq and CD-seq in humans (**k**) and mice (**l**), respectively.

When applied to HEK293T cells, CD-seq achieved >21-fold enrichment of D-box– containing reads compared with conventional small RNA sequencing (sRNA-seq, **Fig. 2b**). Consistently, CD-seq libraries showed markedly higher and more threshold-robust cdRNA detection rates than sRNA-seq (**Fig. 2c**). CD-seq also accurately recovered nearly all annotated cdRNAs (96%, **Extended Data Fig. 3c**) at single-nucleotide resolution (**Fig. 2d, e**), as illustrated by the faithful determination of the 5’ and 3’ ends of ten cdRNAs located within the introns of the GAS5 gene (**Extended Data Fig. 3d**). Importantly, CD-seq showed high reproducibility across replicates and cell types (**Extended Data Fig. 3e, f**; Pearson correlations: CD2-seq, 0.99 and CD5-seq, 0.99), and faithfully captured the expression landscape of known cdRNAs in both human and mouse cell lines (**Extended Data Fig. 3g-o**). Collectively, these results demonstrate that our CD-seq approach not only shows high specificity and accuracy in enriching known cdRNAs, but also offers superior sensitivity and quantitative reliability for low-abundance structured RNAs containing box D motifs, and can capture their full-length sequences.

We next utilized the cdSeeker software to analyze all newly identified Transcription Units (TUs) from CD-seq data. Compared with sRNA-seq, CD-seq uncovered more than 23 times as many klRNAs in the same HEK293T cells (**Fig. 2f**). As with known cdRNAs, the 5’ and 3’ ends of these newly identified klRNAs were resolved at single-nucleotide resolution **(Fig. 2g, h)**. In total, our analysis uncovered 7,023 and 2,207 klRNAs in humans and mice, respectively (**Fig. 2i, j, and Supplementary Table 2 and 4**). Notably, 93.6% of human and 84.9% of mouse klRNAs identified by CD-seq were independently supported by 15.5K RIP-PEN-seq datasets **(Fig. 2k, l)**, providing orthogonal evidence for their association with 15.5K.

We next employed independent experimental methods to further evaluate the accuracy of our CD-seq method. Specifically, we amplified a subset of the newly discovered klRNAs and ktRNAs in various cell lines and tissues using the Poly(T) Adaptor RT-PCR method ^13^ (**Extended Data Fig. 4a**). As expected, the amplicon lengths corresponded precisely to those of the klRNAs and ktRNAs identified by CD-seq (**Extended Data Fig. 4b, c)**. Sanger sequencing confirmed that the sequences of amplicons were also in line with the sequences identified from CD-seq (**Extended Data Fig. 5a, b)**. Together, CD-seq provides orthogonal validation of klRNAs and demonstrates that introns encode a vast and previously hidden landscape of structured RNAs across mammals.

### Introns encode a vast repertoire of compact orphan klRNAs

To define the global landscape of klRNAs across human tissues and cells, we applied cdSeeker software to sequencing datasets generated in this study together with 234 ENCODE sRNA-seq datasets ^14^. This analysis identified 15,213 novel klRNAs (15,584 genomic loci) and 5,659 ktRNAs (spanning 5,817 genomic loci), revealing a previously unrecognized class of compact structured RNAs in the human genome **(Fig. 3a, and Supplementary Table 2)**. Unlike canonical cdRNAs, which typically exceed 60 nucleotides in length and contain extended guide regions, klRNAs are remarkably compact structured RNAs, ranging from 18 nt to approximately 500 nt with a median length of only 36 nt (**Extended Data Fig. 6a**). Strikingly, 85% of klRNA loci mapped to intronic regions **(Fig. 3b)**, collectively distributed across more than 7,000 different host genes (**Extended Data Fig. 6b**). Most klRNA-containing introns encoded a single klRNA locus **(Fig. 3c)**, whereas longer introns tended to encode more klRNAs **(Extended Data Fig. 6c)**, revealing a widespread but nonrandom organization of klRNAs within intronic sequence space.

**Figure 3.**
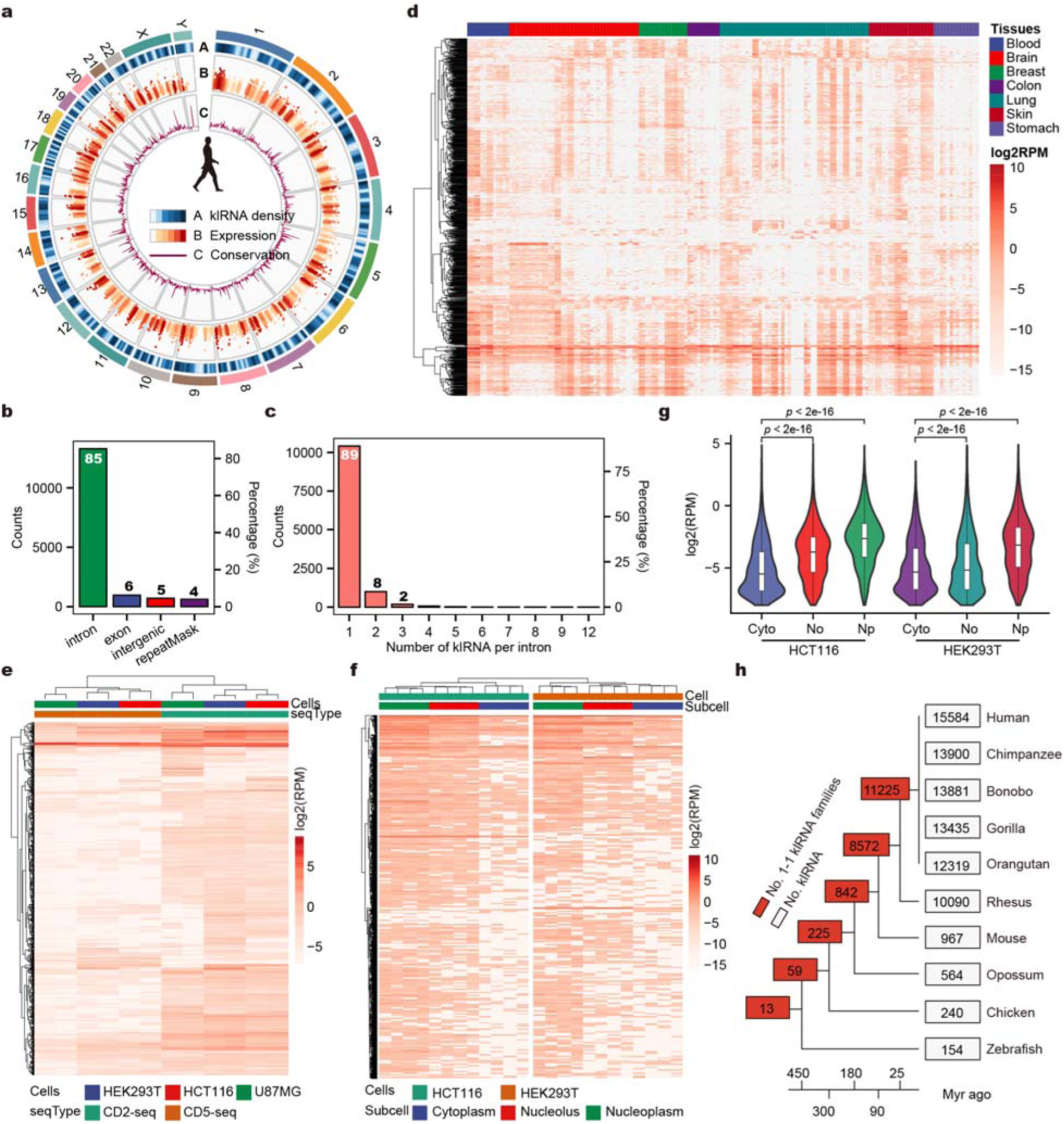
klRNAs comprise an evolutionarily diverse class of orphan intronic structured RNAs. **a,** Circos plot illustrating the genomic distribution, expression and evolutionary conservation of human klRNAs. RPM: reads per million reads. The plot legend is shown in the lower panel. **b,** Numbers and percentage of human klRNAs in different genomic annotation categories. **c,** The count and percentage of introns encoding varying quantities of klRNAs. **d,** Expression landscapes of klRNAs across human tissues datasets. The expression levels of klRNAs in cells were categorized into corresponding tissues. Cells/tissues that have at least 10 datasets are retained. **e,** Heatmap of klRNAs expression in CD-seq libraries. **f,** Expression profiles of klRNAs in different subcellular regions in HCT116 and HEK293T cells. **g,** Violin plots displaying the expression intensity distribution of klRNAs in different subcellular regions in HCT116 and HEK293T cells. The box plots indicate minimum value, first quartile, median, third quartile and maximum value. RPM: reads per million reads. The P values of the differences in distance between the two categories were determined by the Mann-Whitney-Wilcoxon test. Cyto, Cytoplasm; No, Nucleolus; Np, Nucleoplasm. **h,** Simplified phylogenetic trees of human klRNAs. Internal branches and roots, numbers of 1-1 orthologous klRNA families for the indicated species. Tree tips, klRNA numbers for each species. Myr, million years.

Target prediction further revealed that only 0.3% of klRNAs exhibited antisense complementarity to rRNAs or snRNAs, indicating that the overwhelming majority lack the guide potential of canonical cdRNAs and therefore represent orphan klRNAs. Consistent with this distinction, klRNAs differed markedly from canonical cdRNAs in RNA architecture, genomic organization and expression properties **(Extended Data Fig. 6d–j)**. Thus, klRNAs constitute a molecularly distinct and unexpectedly abundant class of intron-encoded orphan structured ncRNAs.

Importantly, independent analysis of mouse 15.5K RIP-PEN-seq and CD-seq datasets identified 3,541 klRNAs together with 1,470 canonical ktRNAs across 5,493 genomic loci, independently establishing the existence of klRNAs in a second mammalian species (**Extended Data Fig. 7a, b, and Supplementary Table 4)**. Consistent with human data, mouse klRNAs exhibit distinct genomic organization and expression properties compared with canonical cdRNAs (**Extended Data Fig. 7c-n)**, supporting klRNAs as a conserved RNA class with shared architectural principles across mammals.

To determine whether klRNAs adopt canonical K-loop folding principles or utilize distinct structural solutions, we analyzed conserved structural features predicted from K-loop folding rules. Structural analysis revealed a strong preference for U pairing at the 3b:3n position (>93%) **(Extended Data Fig. 8a, b)**, consistent with N3 conformations that exhibit reduced metal ion-dependent folding stability ^15–18^. In contrast, only a minority of klRNAs (13.8%, 2,154/15,584) contained canonical stabilizing -1b:-1n base pairs **(Extended Data Fig. 8c)**. Similar structural patterns were observed in mouse klRNAs **(Extended Data Fig. 8d, e)**, indicating that these features are conserved across mammals. Because RNA modifications can influence RNA folding, we further examined m^6^A modifications at structurally relevant positions. Only a small fraction of klRNAs carried m^6^A marks at these positions ^24, 25^ (**Extended Data Fig. 8f, g**), suggesting that klRNA folding is primarily shaped by intrinsic sequence features and RNA-binding proteins rather than a uniform modification-dependent mechanism. Together, these findings indicate that klRNAs adopt K-loop conformations through distinct structural solutions rather than the canonical stabilizing interactions characteristic of cdRNAs.

Transcriptome-wide analyses across diverse cell types and tissues, together with subcellular fractionation, revealed that klRNAs are broadly expressed yet frequently exhibit tissue- and cell-type-specific expression patterns **(Fig. 3d, e, and Extended Data Fig. 8h)**, with predominant nuclear localization **(Fig. 3f, g and Supplementary Table 2)**. RT-qPCR validation across multiple human and mouse tissues confirmed these expression patterns (**Extended Data Fig. 8i, j)**, including highly restricted and ubiquitous subsets (**Extended Data Fig. 9a, b),** confirming that these orphan klRNAs are endogenously expressed in mammalian tissues, with subsets displaying highly restricted or ubiquitous expression patterns. Evolutionary reconstruction further revealed that klRNAs span a wide range of phylogenetic ages, with more than half of klRNA families being primate-specific and a subset predating vertebrate divergence, indicating both rapid evolutionary turnover and long-term conservation **(Fig. 3h)**.

Together, these results establish klRNAs as a widespread class of compact orphan structured ncRNAs encoded within mammalian introns. Their distinct architectures, dynamic expression programs, and diverse evolutionary histories reveal a previously unrecognized layer of RNA-based information embedded within mammalian intronic sequence space.

### The 15.5K protein is required for klRNA biogenesis

Given the direct recognition of klRNAs by 15.5K through their K-loop motifs (**Fig. 1e**), we hypothesized that 15.5K is required for klRNA biogenesis. To address this, we analyzed our previously published PEN-seq datasets generated from two independent 15.5K knockdown cell lines **(Extended Data Fig. 10a)** ^10^. Depletion of 15.5K caused a global reduction in klRNA abundance across both datasets **(Fig. 4a–d and Supplementary Table 5)**, paralleling the reduction observed for canonical cdRNAs **(Extended Data Fig. 10b).** More than 65% of detected klRNAs decreased upon 15.5K depletion **(Fig. 4e)**, and 71% (457/642) of klRNAs quantified in both knockdown cell lines showed concordant reductions **(Fig. 4f)**, indicating a broad dependence of klRNA accumulation on 15.5K.

**Figure 4.**
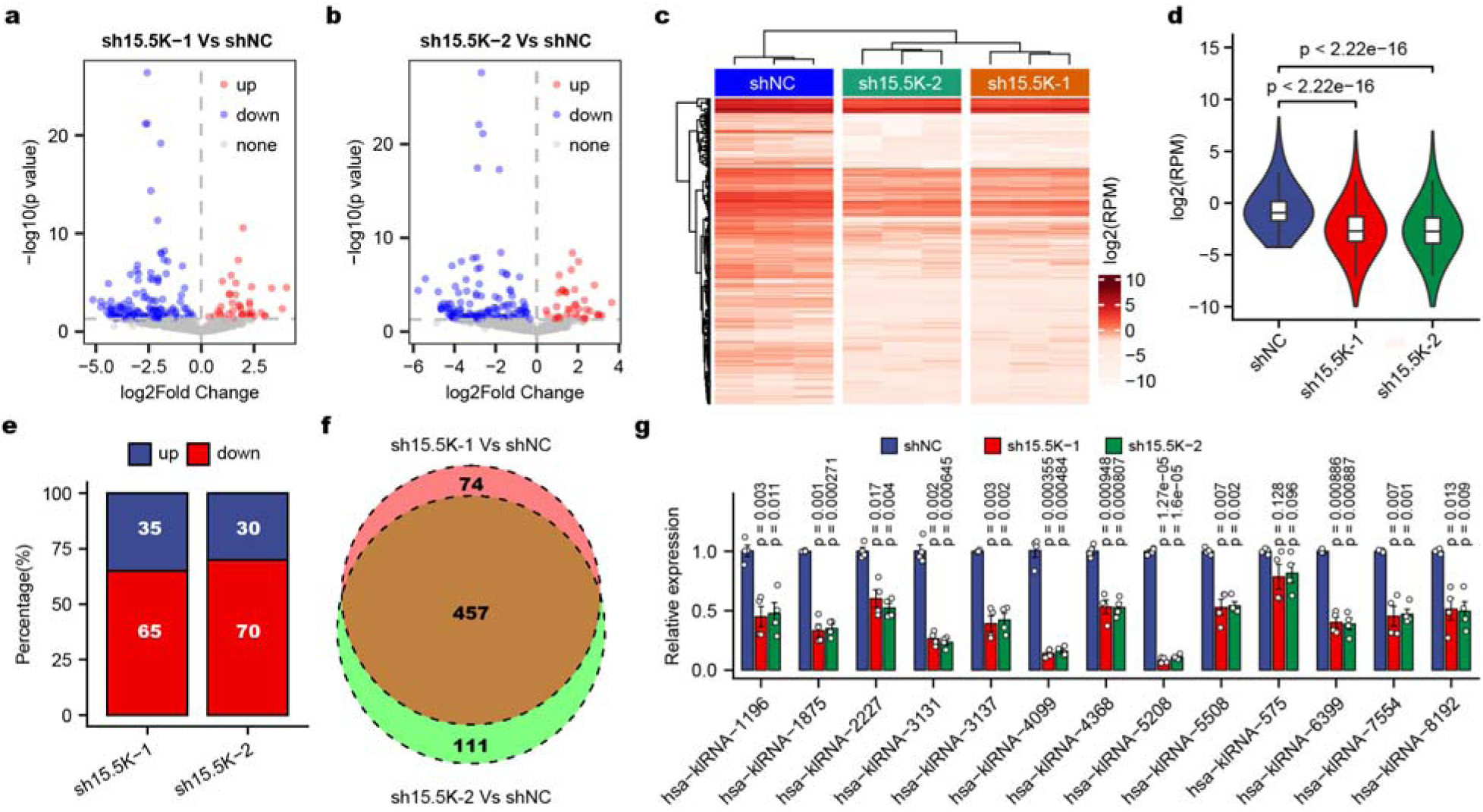
The 15.5K is broadly required for klRNA biogenesis. **a, b,** Differential expression (fold change □≥□ 1.5; P value □<□ 0.05) of klRNAs following 15.5K depletion. **c,** Heatmap displaying expression profiles of klRNAs in 15.5K knockdown and negative control HEK293T cells. RPM: reads per million reads. **d,** Violin plots quantifying the overall expression of klRNAs in 15.5K knockdown and negative control HEK293T cells. Box plots show minimum value, first quartile, median, third quartile and maximum value. RPM: reads per million reads. The P values of the differences in distance between the two categories were determined by the Mann-Whitney-Wilcoxon test. **e,** Percentage of klRNAs dysregulated in 15.5K knockdown cells. **f,** Venn diagram shows the number of downregulated klRNAs between sh15.5K-1 and sh15.5K-2 cells. **g,** qPCR validation of klRNA differential expression upon 15.5K depletion. Data are mean ± s.e.m (n = 4, biological replicates), two-tailed and paired t-test.

To validate these transcriptome-wide observations, we further performed quantitative PCR (qPCR) analysis for a randomly selected subset of klRNAs in 15.5K knockdown cells. Consistent with the 15.5K knockdown PEN-seq data, the majority of the selected klRNAs displayed downregulation in 15.5K knockdown cells (**Fig. 4g**). Together, these findings establish 15.5K as a key determinant of klRNA accumulation, supporting an essential role for 15.5K in klRNA biogenesis.

### klRNAs autoregulate splicing of their host introns through a K-loop-dependent mechanism

Given that klRNAs are predominantly encoded within introns and directly associate with 15.5K, a protein with an established role in spliceosome assembly, we asked whether klRNAs might influence the processing of their host introns. We first examined changes in intron retention following 15.5K depletion using poly(A)+ RNA-seq in HEK293T cells. Consistent with the established role of 15.5K in spliceosome assembly ^10^, 15.5K depletion globally increased intron retention **(Fig. 5a, Extended Data Fig. 10c and Supplementary Table 6 and 7)**. Strikingly, however, klRNA-containing introns deviated from this global trend, exhibiting significantly smaller increases in retention than introns lacking klRNAs **(Fig. 5b)**. Moreover, this countervailing effect became progressively stronger with increasing numbers of klRNAs per intron **(Fig. 5c)**, suggesting a dosage-dependent cis-regulatory effect of intronic klRNAs on host intron processing.

**Figure 5.**
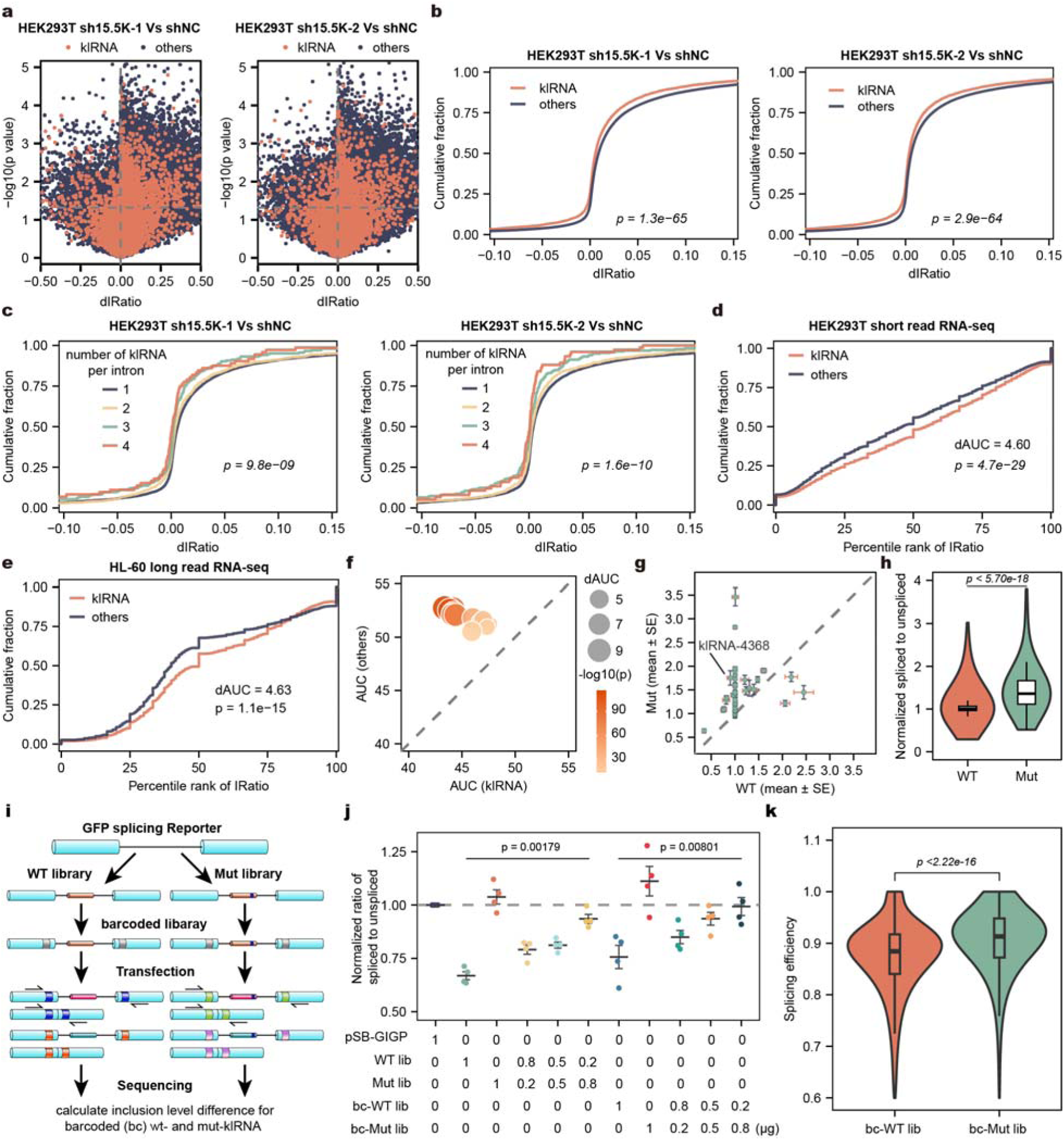
klRNAs autoregulate host intron splicing through K-loop-dependent cis regulation. **a,** Dot plots displaying the intron retention ratio in a representative pairwise analysis of 15.5K knockdown and negative control cells. The orange dots represent klRNA-containing introns and the dark blue dots represent the remaining introns (denoted as others). **b,** Cumulative distribution of differences in intron retention ratios (dIRatio) between klRNA-containing and the remaining introns in 15.5K knockdown and negative control HEK293T cells. The P values on the cumulative plots of inclusion level differences were calculated using a two-sided Mann-Whitney test. **c,** Cumulative fraction of dIRatio for introns harboring different numbers of klRNAs in 15.5K knockdown and negative control HEK293T cells. **d,** Analysis of intron retention ratio for klRNA-containing and klRNA-lacking introns using short-read RNA-seq data from HEK293T cells. The x-axis represents the percentile rank of IRatio, where each intron is ranked according to its retention ratio within the entire dataset, with higher percentiles corresponding to greater intron retention. The delta AUC (dAUC) refers to the difference in the areas under the ECDF curves for intron retention ratios between klRNA-lacking (others) and klRNA-containing introns. The P values on the cumulative plots of inclusion level differences were calculated using a two-sided Mann-Whitney test. **e,** Analysis of intron retention ratio for klRNA-containing and klRNA-lacking introns using long-read RNA-seq data from HL60 cells. The P values on the cumulative plots of inclusion level differences were calculated using a two-sided Mann-Whitney test. **f,** Comparison of the areas under the ECDF curves for intron retention ratios between klRNA-containing introns and other introns, based on short-read and long-read RNA-seq data. The color of each dot indicates the statistical significance determined by two-sided Mann-Whitney tests for cumulative differences in intron inclusion levels. **g,** qPCR analysis of the ratio of spliced to unspliced GFP RNA in HEK293T cells transfected with wild-type (WT) and mutant klRNAs (Mut, mutated from CTGA to CTAG). Horizontal error bars represent the errors of the WT group, and vertical error bars represent those of the Mut group. Data are mean ± s.e.m (n = 4, biological replicates), two-tailed and paired t-test. **h,** Statistical analysis of data from (**g**). Violin plot of normalized spliced to unspliced ratio for wild-type and mutant klRNAs. Box plots show minimum value, first quartile, median, third quartile and maximum value. The P values of the differences in distance between the two categories were determined by the Mann-Whitney-Wilcoxon test. **i,** Schematic of GPS-seq, K-loop mutation library cloning and barcoding strategy. **j,** qPCR analysis of the ratio of spliced to unspliced GFP RNA in cells that were transfected with klRNA-575-based reporters mixed in different ratios. Data are mean ± s.e.m (n = 4, biological replicates). The P values among the indicated categories were determined by the one-way ANOVA. **k,** GPS-seq analysis showing increased splicing following disruption of the K-loop structure. The P values among the indicated categories were determined by the Mann-Whitney-Wilcoxon test.

We next asked whether the association between klRNAs and host-intron processing was evident across independent transcriptomic datasets. Analyses of both short-read RNA-seq and PacBio long-read sequencing across 15 cell lines consistently revealed lower splicing efficiency for klRNA-containing introns than for introns lacking klRNAs (**Fig. 5d-f)**. Notably, this effect was independent of klRNA abundance or distance to splice site **(Extended Data Fig. 10d-f)**, indicating that klRNA-mediated regulation represents an intrinsic property encoded within intronic sequences rather than a secondary consequence of gene expression level or genomic context.

To directly test whether klRNAs can regulate intron splicing, we constructed GFP-based reporters containing individual klRNAs within their native intronic contexts and compared wild-type constructs with corresponding D-box mutants **(Extended Data Fig. 11a)**. Among 40 klRNAs tested, 26 (65%) significantly suppressed intron splicing relative to their D-box-mutant counterparts **(Fig. 5g, h and Extended Data Fig. 11b)**, revealing regulatory activity across a substantial fraction of tested klRNAs. We further varied the ratio of wild-type and D-box-mutant klRNAs across 80 GFP reporters and observed quantitative changes in splicing efficiency with the proportion of intact klRNA **(Extended Data Fig. 11c)**, supporting a requirement for an intact K-loop motif in klRNA-mediated splicing regulation.

To systematically test whether this regulatory activity is encoded by K-loop architecture rather than the intervening sequence, we developed GFP Splicing Reporter sequencing (GPS-seq), in which intronic regions between C and D boxes were replaced with randomized reporter sequences (**Fig. 5i**). As an initial validation, qPCR analysis of representative reporters, including klRNA-575, showed that D-box mutation increased intron splicing relative to the wild-type construct (**Fig. 5j**), consistent with a repressive role of klRNAs in local splicing. We next applied GPS-seq to barcoded reporter libraries to quantify splicing outcomes at scale. Across thousands of reporter constructs, D-box mutations consistently increased splicing efficiency (**Fig. 5k**), demonstrating that klRNA-mediated splicing inhibition depends on the K-loop structural architecture rather than the intervening sequence composition between C and D boxes.

To test the function of endogenous klRNAs in host-intron splicing, we performed CRISPR-Cas9–mediated deletion of klRNA-575 and klRNA-7554 in HEK293T cells (**Fig. 6a**). Successful genomic deletion was confirmed by PCR (**Extended Data Fig. 11d**). Loss of these klRNAs led to a selective increase in splicing efficiency of their host introns, while leaving distant introns unaffected (**Fig. 6b, c**), supporting a local *cis*-regulatory function. To directly assess the structural requirement of endogenous klRNAs, we introduced precise D-box mutations in klRNA-575 using prime editing (CTGA→CTAG) **(Fig. 6d**). Sanger sequencing confirmed generation of homozygous mutant clones (**Extended Data Fig. 11e**). Consistent with reporter-based assays, disruption of the D box resulted in increased splicing efficiency of the corresponding intron, while splicing of adjacent introns within the same gene remained unchanged (**Fig. 6e**). Notably, prime editing–induced disruption of the K-loop motif also reduced local association of 15.5K with the affected intronic region (**Fig. 6f**), linking structural integrity of klRNAs to protein recruitment *in vivo*.

**Figure 6.**
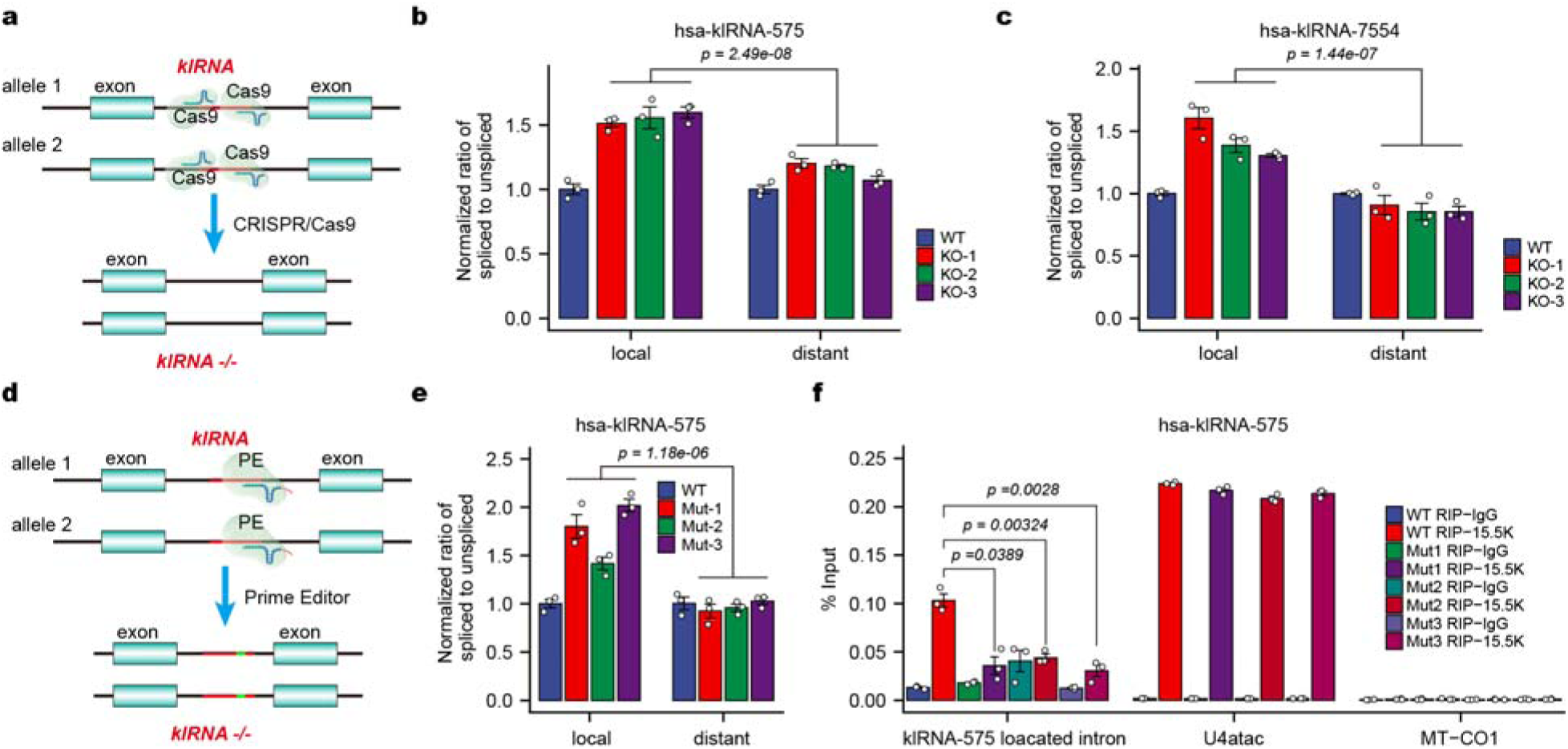
Endogenous klRNAs autoregulate host intron splicing in vivo. **a,** CRISPR–Cas9 strategy for deletion of endogenous klRNA loci. **b, c,** Quantification of local and distal intron splicing following deletion of klRNA-575 (**b**) and klRNA-7554 (**c**). Data are mean ± s.e.m (n = 3, biological replicates). The P values among the indicated categories were determined by the one-way ANOVA. **d,** Prime-editing strategy for disruption of the klRNA D-box. **e,** Effects of K-loop disruption on host intron splicing in HEK293T cells expressing wild□ type or D□ box mutated klRNA□575. Data are mean ± s.e.m (n = 3, biological replicates), two-tailed and paired t-test. **f,** RIP–qPCR analysis of endogenous 15.5K association with wild-type and D-box mutant klRNA loci. U4atac and Mt-CO1 were kept as positive and negative controls, respectively. Data are mean ± s.e.m (n = 3, biological replicates), two-tailed and paired t-test.

Together, these functional and genetic perturbations establish that klRNAs suppress host intron excision *in cis* through a K-loop- and 15.5K-dependent mechanism. Thus, klRNAs define a widespread mechanism by which introns autoregulate their own splicing (**Extended Data Fig. 11f**).

## Discussion

In this study, we identify a widespread new class of intron-encoded K-loop RNAs (klRNAs) that reveal a previously unrecognized layer of RNA structure–encoded regulation within mammalian introns. Using two orthogonal sequencing strategies, eRIP-PEN-seq and CD-seq, we demonstrate that introns represent a vast and previously underexplored source of structured RNAs specifically recognized by the conserved RNA-binding protein 15.5K. Despite their compact size and generally low abundance, klRNAs are broadly distributed throughout mammalian genomes, exhibit regulated expression patterns across tissues and cell types, and function as cis-regulatory elements that control the processing of their host introns. These findings reveal that introns can encode structured regulatory RNAs that govern their own fate.

Comprehensive analyses demonstrate that klRNAs constitute a distinct class of structured RNAs fundamentally different from canonical cdRNAs. Compared with canonical cdRNAs, klRNAs represent a repertoire more than 50-fold larger than the known canonical cdRNA repertoire ^19^, are remarkably compact (median length of 36 nt), and form K-loop structures through stereotypically positioned terminal C/D motifs rather than internal C′/D′ motifs of canonical archaeal cdRNAs. Moreover, klRNAs overwhelmingly lack antisense complementarity to rRNAs or snRNAs, defining them as orphan structured RNAs rather than conventional modification guides. Their distinct structural architectures, expression programs and evolutionary histories further distinguish them from canonical cdRNAs. Together, these features establish klRNAs as an unexpectedly extensive class of intron-encoded orphan structured RNAs in mammalian genomes.

The vast repertoire of klRNAs uncovered in this study likely reflects both their widespread genomic distribution and their previous inaccessibility to conventional RNA profiling approaches. Standard small RNA sequencing is biased toward abundant or predefined short RNA classes (e.g., miRNAs and piRNAs), while highly abundant housekeeping ncRNAs, including rRNAs, tRNAs and snRNAs, can further obscure low-abundance structured transcripts. By contrast, eRIP-PEN-seq selectively enriches 15.5K-associated RNAs through depletion of dominant ncRNAs, whereas CD-seq independently captures full-length D-box–containing RNAs with high sensitivity and precise end resolution. The convergence of these orthogonal approaches provides independent support for the prevalence of klRNAs while overcoming key limitations of conventional RNA profiling. Given the limited cell types and tissues examined here, the mammalian klRNA repertoire is likely substantially larger than currently defined, particularly across diverse developmental, physiological and disease states.

Our findings also expand the structural landscape of K-loop recognition. In canonical archaeal cdRNAs, K-loops occur at internal C′/D′ motifs, whereas klRNAs deploy K-loop architecture at stereotypically positioned terminal C/D motifs. Despite lacking canonical C-stems, these RNAs adopt K-loop conformations *in vivo* and are directly recognized by 15.5K. Their heterogeneous terminal pairing patterns and limited dependence on RNA modifications further suggest that klRNAs achieve compatible K-loop conformations through multiple structural solutions. Thus, K-loop architecture can function as a terminal structural principle defining an distinct class of mammalian RNAs.

Most importantly, klRNAs reveal an unexpected mechanism by which introns regulate their own processing. Intron retention contributes broadly to gene regulation in development, differentiation and disease ^20^, but whether introns encode discrete RNA structures that determine their own splicing has remained poorly understood. Intron-retention analyses, reporter assays, high-throughput functional screening and endogenous genetic perturbations consistently show that klRNAs suppress host intron excision through a K-loop-dependent mechanism. Precise disruption of the endogenous K-loop further increases host-intron splicing while reducing local 15.5K association, directly linking RNA structure, protein recruitment and splicing regulation. We propose that formation of the klRNA–15.5K complex locally influences spliceosomal assembly or progression, thereby delaying intron removal. In this model, an intron is not simply a substrate for splicing but encodes a structured RNA element that feeds back on its own excision. How klRNA–15.5K complexes interface with the spliceosome, and whether this regulatory axis is dynamically controlled during development or disease, will be important questions for future investigation.

More broadly, the discovery of thousands of klRNAs expands the functional landscape of mammalian intronic sequence space. Rather than serving solely as sequences to be removed from pre-mRNAs, introns can encode autonomous structural information that regulates their own processing. klRNAs therefore establish RNA structure–encoded intron autoregulation as a widespread regulatory principle and reveal RNA tertiary structure as an additional layer through which intronic sequences encode biological information.

## Methods

### Cell culture

HEK293T, HCT116, U-87 MG, C2C12, and Hepa1-6 cell lines were sourced from the Cell Bank of China Science Academy in Shanghai, China. These cell lines were cultured in DMEM (Gibco) supplemented with 10% fetal bovine serum (FBS, Gibco) and 1% penicillin-streptomycin solution (Gibco) at 37 in a 5% CO2 environment. The cells were free of mycoplasma contamination based on the MycoBlue Mycoplasma Detector (Vazyme), and STR profiling was used to authenticate the cell lines (Cellcook Biotech Co., Ltd, Guangzhou).

### Construction of plasmids

Recombinant vectors encoding human klRNAs and mutants (extending by 30 bp on each side flanking the klRNAs) were amplified using the overlap extension PCR strategy and then cloned using the *Bsa* I restriction site in the pSB-GIGP ^10, 21^. To achieve klRNA deletion, dual-sgRNAs targeting the 5’ and 3’ regions flanking the klRNAs locus were subcloned into PX459 (Addgene #62988) by restriction cloning. The pegRNAs (including sgRNA, RTT and PBS sequences) targeting klRNAs were subcloned into Prime Editor pCMV-PE2-Puro ^10^. Sequences of all primers or oligonucleotides are listed in **Supplementary Table 8**.

### Generation of klRNA knockout and prime edited cell lines

For generation of klRNA knockout or D box-mutated HEK293T cell lines, the targeting vectors PX459-sgklRNA or pCMV-PE2-Puro-pegRNA were transfected into HEK293T cells with Superluminal (MIKX). Twenty-four hours post-transfection, 4 μg/ml puromycin was added to the culture medium. Forty-eight hours after antibiotic selection, individual colonies were picked via 2-fold serial dilution assays. Clones were screened by PCR for deletion of the klRNA locus. For validation of target DNA editing, genomic DNA was isolated from the engineered cells, and the target loci were amplified with the corresponding primers (klRNA-VF1 and klRNA-VR1). The amplified DNA sequence was cloned into a Zero blunt TOPO vector (Vazyme) according to manufacturer instructions and subjected to Sanger sequencing analysis to determine biallelic gene alterations. The primers used for knockout cell validation are listed in **Supplementary Table 8**.

### eRIP-PEN-seq library preparation

eRIP-PEN-seq was conducted following the protocol of RIP-PEN-seq ^10^ with some modifications. To enhance the read coverage on klRNAs, we designed specific probes for RNase H-based high-abundance snoRNA subtraction, in addition to depleting rRNA and snRNA. The high-abundance snoRNAs were selected based on the top 1000 reads in the 15.5K RIP-PEN-seq datasets. Details of the adapters, primers, and probes can be found in **Supplementary Table 8**.

### Construction of CD-seq and small RNA-seq libraries

CD-seq employs a D box-anchored ligation approach to generate D box-specific ligation products. Briefly, a recognition site for the type II restriction endonuclease *Btg*ZI is introduced at the 3’-terminus of RNAs via pre-adenylated 3’-adaptor ligation. Following digestion of the remaining 3’-adaptor, a 5’-adaptor is ligated to capture information at the 5’ extremity of RNA. The cDNAs are then reverse transcribed using a specific primer complementary to the 3’-adaptor. Subsequently, the cDNAs are amplified by low-cycle PCR and cleaved by *Btg*ZI at a sequence 10 and 14 nucleotides downstream (3’) from its recognition site. This cleavage removes the sequence beyond the motif site at the 3’-terminus and creates a motif-containing sticky end with a four-nucleotide overhang. DNAs terminating with the D box sequence are then preferentially ligated to an adaptor with a complementary four-nucleotide (5’-CTGA) overhang, thereby increasing the detectability of klRNAs. Moreover, we used a D box-anchored PCR amplification strategy to generate the library, which represents the core step for klRNA enrichment and provides a format for downstream bridge amplification and sequencing. Given klRNAs generally leave two or five nucleotides after the D-box (**Fig. 2a**), we designed CD2-seq and CD5-seq protocols to capture these variants (**Fig. 2a and Extended Data Fig. 3a, b)**.

Total RNA was extracted using RNAzol ^22^. Then the total RNA was treated with DNase I (Promega) to remove genomic DNA and purified using RNA Clean & Concentrator-5 (Zymoresearch). After ligation of 3’ RNA adaptor (CD2RA3 or CD5RA3) to 1 μg total RNA with T4 RNA ligase 2 truncated KQ (NEB) in 1× T4 RNA ligase reaction buffer supplemented with 12.5% PEG 8000 at 16 °C for 18 h, the excessive adaptors were digested with 100 U 5’ Deadenylase (NEB) at 30 °C for 1 h, incubated with 2 μg E. coli Single-strand DNA-binding protein (Promega) on ice for 30 min and subjected to ssDNA digestion with 60 U of RecJf (NEB) at 37 °C for another hour ^23, 24^. The ligated RNA was ligated to a 5’ RNA adaptor (RA5) using T4 RNA Ligase 1 (NEB) in 1× T4 RNA ligase reaction buffer supplemented with 1 mM ATP at 16 °C for 18 h. The ligated RNA was column-purified by RNA Clean & Concentrator-5 (Zymo Research) and subjected to reverse transcription with specific primers (CD2RTP or CD5RTP) and SuperScript IV Reverse Transcriptase (Thermo Fisher). For cDNA purification, *Exo* I (NEB) was used to digest excess RT primers at 37 □ for 15 min. Then, 7 μl 1 M NaOH and 5 μl 0.5 M EDTA per 20 μl reaction volume was added to remove RNA templates at 70□ for 12 min, after which clean-up of the cDNA was performed with Oligo Clean & Concentrator (Zymo research). Following low-cycle (10 cycles) PCR amplification with DA3sP and CD2RTP or CD5RTP primers, the PCR products were purified with DNA Clean & Concentrator-5 kit and subjected to *Btg*Z I digestion and PAGE-gel purification. The CTGA-anchored adaptors were prepared by annealing CDDA3F and CDDA3R oligos. The annealing was conducted as follows: 2 μl corresponding oligos at 100 μM each were combined in 1 μl of 10× NEB Buffer 2 and 5 μl Nuclease-free Water, incubated at 95 °C for 4 min and then slowly (0.05 □/s) cooled to 25 °C. Then, the *Btg*Z I-digested DNA was ligated to CTGA-anchored adaptors using 1200U T4 DNA Ligase (NEB) overnight at 16 °C in 1 × buffer and a 50-μl reaction volume, purified with DNA Clean & Concentrator-5 kit. Purified DNA was subjected to CTGA-anchored PCR amplification with RP1 and CD-Index primers. Amplified DNA was purified on a 4% low-melting agarose gel with a size selection of 150-700 bp. For small RNA-seq library, the DNase I-treated RNA was subjected to library generation using 3’ RNA adaptor (RA3), 5’ RNA adaptor (RA5), RT primer (RTP) and PCR primers RP1 (forward primer) and RPI1-4 (reverse primer) as above. And the PCR products were run on a 4% low-melting agarose gel and the size range from 150 bp to 700 bp was recovered by Zymoclean™ Gel DNA recovery Kit. The libraries were sequenced through a 150-bp paired-end (PE) run on the Illumina HiSeq X platform (Illumina) at Annoroad Gene Technology Company. The adaptors and primers are listed in **Supplementary Table 8**.

### Expression and purification of 15.5K protein

15.5K protein was expressed as a C-terminal His (6×) tags fusion in Escherichia coli Rosetta (DE3) cells. *E. coli* were grown to OD600 of 1.0 and induced with 0.2 mM IPTG at 16°C overnight. Pelleted cells were resuspended in lysis buffer (500 mM NaCl, 20 mM Tris-HCl pH8.0, 10%(v/v) glycerol, 1 mM DTT, 30 mM imidazole, 0.4 mM PMSF, 0.5 ng/mL pepstatinA). After sonication, lysates were spun at 13,200 × g, 4°C for 30 min. The solution was then applied to a HisTrap column equilibrated in binding buffer (500 mM NaCl, 20 mM Tris-HCl pH8.0, 10%(v/v) glycerol, 1 mM DTT, 30 mM imidazole). Proteins were eluted in elution buffer (500 mM NaCl, 20 mM Tris-HCl pH8.0, 10%(v/v) glycerol, 1 mM DTT, 300 mM imidazole), and then dialyzed in 20 mM Tris–HCl pH 8.0, 50 mM NaCl, 1mM DTT at 4°C overnight. In order to remove contaminating RNA and other impurities, the proteins were applied to a heparin column (HiTrap Heparin HP, 5mL) in 20 mM Tris–HCl pH8.0, 1mM DTT, 10%(v/v) glycerol with a 50–1,000 mM NaCl gradient. The fractions were analyzed by SDS-PAGE, pooled, concentrated, filtered, flash frozen in liquid nitrogen, and stored at -80°C.

### RNA electrophoretic mobility shift assays (REMSA)

REMSA was performed as previously described with some modifications ^10^. In brief, Cy5-labeled RNA probes were denatured in nuclease-free water by incubation at 95□ for 2 minutes, followed by quick cooling on ice. Recombinant 15.5K protein was diluted to various concentrations of 0 μM, 2 μM, 6 μM, 10 μM, and 14 μM in protein dilution buffer (containing 50 mM HEPES, pH 7.5, 150 mM NaCl, 0.5 mM EDTA, and 0.005% NP-40), supplemented with 1 mM DTT. For each reaction, 1 μl of RNA probes was used, resulting in a final concentration of 8 nM, along with 1 μl of 15.5K protein (final concentrations of 0 nM, 100 nM, 300 nM, 500 nM, and 700 nM). This mixture was then incubated in 5 μl of 4× REMSA Binding Buffer (comprising 80 mM HEPES, pH 7.5, 500 mM KCl, 20% Glycerol, 0.4% TritonX-100, 6 mM MgCl2, and 4 mM DTT), with the addition of 2 μg of yeast tRNA (ThermoFisher) and 1 μl of Murine RNase inhibitor (Vazyme). The incubation was carried out at 30□ for 30 minutes, and after adding 5 μl of 5× loading buffer (consisting of 50 mM HEPES, pH 7.5, 80% glycerol, and 0.25% bromophenol blue), the samples were separated by 6% native PAGE. The fluorescence signal was detected using Odyssey Imaging Systems (Licor) and subsequently quantified via Image Studio (Licor). The dissociation constant (Kd) was then calculated through nonlinear curve fitting. The equation used for the fitting was Y = Bmax × X / (Kd + X), where Y represents the ratio of [RNA–protein] / ([freeRNA] + [RNA–protein]), X stands for the input protein concentration, and Bmax was set to 1.

### RNA isolation and quantitative reverse transcriptase PCR (qPCR)

Total RNA was isolated from cells with RNAzol ^22^ and then used for cDNA preparation with the HiScript II Q RT SuperMix for qPCR (+gDNA wiper) kit (Vazyme). qPCR was performed using TB Green Premix Ex Taq II (Tli RNaseH Plus) (Takara) or ChamQ Universal SYBR qPCR Master Mix (Vazyme) with gene-specific primers. The ΔΔCt method versus GAPDH for mRNA or U6 for small RNA was applied to calculate expression differences. For splicing efficiency analysis, the ratio of spliced to unspliced pre-mRNA was determined by qPCR. The primer sequences are listed in **Supplementary Table 8**.

### Poly(T) adaptor RT-PCR-based klRNA validation and quantification

RNA samples from 10 normal human tissues were purchased from Clontech (Mountain View, CA). The poly(T) adaptor RT-PCR analysis of the RNAs was carried out according to the original protocol with minor modifications ^13^. Briefly, total RNAs from cultured cells or tissues were extracted using RNAzol ^22^, followed by RQ1 RNase-free DNase treatment (Promega). Prior to poly(A) tailing and reverse transcription, DNase Stop solution was added to each reaction. The DNase-treated RNA was extended through a poly(A) tailing reaction (E. coli Poly(A) Polymerase, NEB) and then reverse transcribed into cDNA using a poly(T) adaptor with HiScript ® II Reverse Transcriptase (Vazyme). A gene-specific forward primer and a universal poly(T) adaptor reverse primer were used to amplify and validate the klRNAs. The comparative Ct Method (ΔΔCT Method) was used to determine the expression levels of the genes. For agarose gel electrophoresis, cDNA was amplified with the forward and reverse primers by using Premix Ex Taq™ Hot Start Version (Takara) and then resolved in 2.5% agarose gels. For poly(T) adaptor RT-qPCR, we used TB Green Premix Ex Taq II (Tli RNaseH Plus) (Takara) and the forward and reverse primers following the provided instructions. U6 was employed as an endogenous control. For Sanger sequencing validation, the amplified products were subcloned into Zero blunt TOPO vector (Vazyme) according to manufacturer instructions and subjected to Sanger sequencing. The primers used for poly(T) adaptor RT-PCR or qPCR are listed in **Supplementary Table 8**.

### GFP Splicing reporter assay and sequencing (GPS-seq)

To systematically address the impact of the sequence and K-loop structure of klRNA to local intron splicing, a minigene reporter based on hsa-klRNA-575 was constructed in the pSB-GIGP backbone. Synthetic DNA (wt-klRNA-575, mut-klRNA-575) pieces, containing wild-type and D box-mutated K-loop structure and the corresponding flanking sequences were amplified by Amp-F/R, respectively. The amplified DNA sequences were Gibson assembled into *Bsa* I-digested pSB-GIGP and electroporated into *E.coli* DH5α Electro-Cells (Takara), resulting in non-barcoded library plasmids (pWT575 and pMut575).

To generate barcoded libraries, an intermediate vector (pSB-med) was generated as follows: Gmed-CF1/CR1 and Gmed-CF2/CR2 primer pairs were used to amplify the corresponding GFP fragments and then the amplified DNA sequences were Gibson assembled into *Bgl* II and *Apa* I digested pSB-GIGP. The barcoded DNA fragments were amplified from non-barcoded library plasmids by using R9index-CF/CR. Afterwards, the amplified products were Gibson assembled into *EcoR* V-digested pSB-med and electroporated into *E.coli* DH5α Electro-Cells (Takara), resulting in barcoded libraries (pbcWT575 and pbcMut575). The barcoded libraries were then transiently transfected into HEK293T cells by Superluminal (MIKX). 48 h post transfection, the RNA was isolated and subjected to library construction.

Total RNA extracted from barcoded libraries-transfected cells, was first DNase treated using RQ1 RNase-Free DNase (Promega) and was then subjected to reverse transcription using SuperScript IV Reverse Transcriptase with specific primer (GFP-RT). Following pre-PCR amplification with GFP-libF/R primers by Q5 Master Mix (NEB), the PCR products were purified using 0.9× VAHTS DNA Clean Beads (Vazyme). Then the retrieved DNA was PCR amplified with Illumina index primers and the libraries were purified using 0.8× VAHTS DNA Clean Beads (Vazyme) and sequenced on the Illumina NovaSeq platform with 150-bp paired-end reads at Annoroad Gene Technology Company. The oligos and primers are listed in **Supplementary Table 8**.

### Mice

All BALB/c mice (Guangdong Medical Lab Animal Center) were bred and maintained in the animal facility of the Laboratory Animal Center of Sun Yat-sen University under specific pathogen-free conditions, in plastic cages and were provided regular chow and water ad libitum. All aspects of animal care and the experimental protocols were approved by the Animal Ethics Committee of Sun Yat-sen University. Adult mice were dissected and tissue samples were collected and named according to The Anatomy of the Laboratory Mouse (http://www.informatics.jax.org/cookbook/index.shtml). Tissue samples were collected directly in ice-cold RNAlater solution (25 mM Sodium Citrate, 20 mM EDTA, 5.3 M ammonium sulfate, pH 5.2). Then RNA was extracted from mouse tissues stored in RNAlater solution with RNAzol.

### Computational analysis of eRIP-PEN-seq, RIP-PEN-seq, CD-seq and sRNA-seq data Read processing and alignment

Paired 5′- and 3′-end reads from eRIP-PEN-seq, RIP-PEN-seq, CD-seq, and sRNA-seq libraries were trimmed with Cutadapt (v4.4) ^25^ to remove adaptors and were then mapped to the human (hg38) and mouse genome (mm10) using the STAR program ^26^ with the following parameters: ‘--alignEndsType EndToEnd --outFilterMultimapScoreRange 0 -- outFilterMultimapNmax 20 --outFilterMismatchNmax 10 --outFilterMismatchNoverLmax 0.05 - -outFilterScoreMin 0 --outFilterScoreMinOverLread 0 --outFilterMatchNmin 15 -- outFilterMatchNminOverLread 0 --alignIntronMin 1 --alignIntronMax 1 –alignMatesGapMax 1500 --seedSearchStartLmax 15 --seedSearchStartLmaxOverLread 1 --seedSearchLmax 0 -- seedMultimapNmax 20000 --seedPerReadNmax 1000 --seedPerWindowNmax 100 -- seedNoneLociPerWindow 20 --alignSJDBoverhangMin 1000’. Reads mapping to more than twenty locations were discarded and only sequences in which both ends were aligned to the reference genomes were further analyzed.

### Identification and classification of klRNAs and ktRNAs

Overlapping paired-end reads mapped to the genome were first clustered into transcriptional units (TUs). Within each cluster, the genomic positions showing the highest abundance of 5’ and 3’ ends were designated as the transcript start (TSS) and terminal sites (TTS), respectively. Candidate transcripts were reconstructed from paired start and terminal sites occurring within 500 bp.

Candidate transcripts were then analyzed using cdSeeker, implemented based on the cdRNA identification framework of our previously developed snoSeeker algorithm ^11^, to identify terminal C- and D-box motifs. Candidates with a cdSeeker score of ≥11.5 bits were retained and further required to satisfy the following criteria: (1) a D-box motif (CUGA) was positioned proximal to the 3′ terminus; (2) a C-box motif (RUGAUGA) was located within 7 nt of the 5′ terminus; and (3) the candidate was detected in at least two sequencing libraries.

To distinguish K-loop RNAs (klRNAs) from K-turn RNAs (ktRNAs), terminal base-pairing between sequences flanking the C- and D-box motifs was evaluated according to the structural criteria used for K-turn/K-loop classification. Candidates containing a canonical C-stem comprising at least two terminal base pairs were classified as ktRNAs. Candidates containing fewer than two C-stem base pairs were classified as klRNAs, representing terminal C/D motif-containing RNAs lacking the canonical C-stem required for K-turn architecture. Thus, klRNAs were defined by the combination of stereotypically positioned terminal C/D motifs and the absence of a canonical C-stem.

### Annotation of novel klRNAs

The genome sequences of humans (hg38) and mice (mm10) were downloaded from the UCSC Genome Browser site ^27^. Human and mouse gene annotations were acquired from GENCODE ^28^. The repeat elements in RepeatMasker were downloaded from the UCSC Genome Browser site ^27^. Reference annotations of canonical human and mouse cdRNAs were compiled from snoRNA-LBME-db were downloaded from snoRNA-LBME-db ^19^, deepBase ^29^, GENCODE ^28^, snoRNAome ^30, 31^ and Refseq ^32^. All novel cdRNAs were further intersected with canonical gene annotations using bedtools software ^33^ to determine their genomic origins, including intronic, exonic, and intergenic locations..

### Gene Ontology (GO) enrichment analyses of the host genes of klRNAs

GO enrichment was performed using the R package clusterProfiler version 2.2.5 ^34^. The host genes of klRNAs identified from CD-seq or eRIP-PEN-seq data were analyzed using the compareCluster function with the following settings: fun = enrichGO, OrgDb = org.Hs.eg.db, pAdjustMethod = ’BH’, and pvalueCutoff = 0.05.

### Construction of homologous klRNA families

We constructed homologous klRNA families based on DNA sequence similarity. To identify homologous human klRNA families in other species, we used the UCSC liftOver tool ^27^ to search the candidate klRNA from the *Pan troglodytes* reference genome (chimpanzee, panTro4), the *Pan paniscus* reference genome (bonobo, panPan2), the *Gorilla gorilla gorilla* reference genome (gorilla, gorGor4), the *Pongo pygmaeus abelii* reference genome (orangutan, ponAbe3), the *Macaca mulatta* reference genome (rhesus, rheMac8), the *Mus musculus* reference genome (mouse, mm10), the *Monodelphis domestica* reference genome (opossum, monDom5), the *Gallus gallus* reference genome (chicken, galGal6), and the *Danio rerio* reference genome (zebrafish, danRer11), respectively. Then we filtered these candidates with cdSeeker as mentioned above.

### Prediction of klRNA targets

Experimentally annotated targets (2’-O-Methylation sites) were obtained from snoRNABase ^19^. Potential antisense targets of klRNAs were predicted using snoSeeker ^11^. Only predicted interactions with a snoSeeker target score >15.0 were retained for downstream analyses.

### Analysis of RIP-PEN-SHAPE-MaP data

RIP-PEN-SHAPE-MaP sequencing reads were processed by removing adaptor sequences using Cutadapt, merging paired-end reads using FLASH2 ^35^, and collapsing PCR duplicates based on the 6-nt unique molecular identifiers (UMIs) at both the 5′ and 3′ ends using the fastq tool from the bio-playground package. Processed reads were aligned using STAR to a customized reference comprising all identified klRNA and canonical cdRNA sequences together with the hg38 genome, in which the corresponding genomic klRNA and cdRNA loci were masked to minimize ambiguous mapping. STAR was run with the following parameters: -- alignSJoverhangMin 1000 --scoreDelOpen -1 --scoreDelBase -1 --scoreInsOpen -1 -- scoreInsBase -1 --outFilterMultimapScoreRange 1 --outFilterMultimapNmax 20 -- outFilterMismatchNmax 30 --outFilterMismatchNoverLmax 0.3 -- outFilterMismatchNoverReadLmax 1 --outFilterScoreMin 0 --outFilterScoreMinOverLread 0.3 - -outFilterMatchNmin 25 --outFilterMatchNminOverLread 0. Alignments corresponding to individual klRNAs and canonical cdRNAs were extracted and analyzed using shapemapper_mutation_parser, shapemapper_mutation_counter, and make_reactivity_profiles.py from ShapeMapper2 ^36^ to calculate nucleotide-resolution SHAPE reactivity profiles. RNAs were retained for structural analysis only when both control and SHAPE-treated samples achieved a median effective coverage of ≥100 across at least 80% of transcript positions. Nucleotide-resolution SHAPE reactivities were mapped onto predicted RNA secondary structures and visualized using R2easyR ^37^ and R2R ^38^.

### Analysis of GPS-seq data

For each paired-end reads from GPS-seq data, the read1 and read2 are matched against the mode “[5’] ACCACATGAAGCAGCACGAT[9 nt barcode]TTCTTCAAGTCCGCCATGCCCGAAG” and “[5’]GCCCTCGAACTTCACCTCGGCGCGGGTCTTGTAGTTGCCGTCGTCCTTGAAGAAGAT[9 nt barcode]GGTGCGCTCCTGGACGTAGC”. The 9 nt barcodes from the two reads are combined into an 18 nt barcodes that represent the identity of the transcript. Next, if the read 1 further matches the mode “[5’][54 nt]GTTGGTATCAACTCGAGGGTTACAAGACAGGTTTAATGACCGTGGCTAAAAATCCCAAGTGACCTCATGATGG[23 nt internal sequence][3’]”, the paired-end read is considered to support intron retention. While if the read 1 matches the mode “[5’][54 nt]GCTACGTCCAGGAGCGCACC[9 nt barcode]ATCTTCTTCAAGGACGACGGCAACTACAAGACCCGCGCCGAGGTGAAGTTCGAGGGCAGATCGGAAG[3’]”, the paired-end read is considered to support the spliced transcript. To remove the barcodes derived from potential sequencing error, we clustered all barcodes that differ less than 2 nt and keep the barcodes only if they account for at least 95% of all reads belonging to the cluster. The resulting spliced GFP reads were used to match the reporter barcodes with the unspliced GFP reads. Splicing efficiency is defined as the amount of spliced GFP relative to total GFP (spliced GFP + unspliced GFP).

### mRNA-seq differential expression analysis

Gene expression level measured as raw counts was determined by in-house developed software with annotated klRNAs and snoRNAs as references. Differential expression was analyzed using DESeq2 workflow ^39^.

### mRNA-seq differential splicing analysis

Adaptor sequences were trimmed from raw RNA-seq data using Cutadapt. The clean reads were mapped to the reference genome (hg38) using the STAR software (2.7.1a) ^26^ with the genome index built from GENCODE v39 ^28^ annotation and with the following additional parameters: -- alignEndsType EndToEnd --outSAMstrandField intronMotif --outFilterMismatchNmax 5. The genomic coordinates of introns were extracted from GENCODE v39 ^28^ annotation and duplicates were further removed. For each intron, the numbers of reads supported either the spliced isoform or the retained isoform were counted. Reads that skipped the intron and spanned at least 10 bp in both exons were considered to support the spliced isoform, while those that included the intron with at least 10 bp overlapping between the exon side and intron side were considered to support the retained isoform. The read counts were normalized by effective length, defined as the number of possible positions for supporting reads, to estimate the abundance of the spliced isoform and the retained isoform. Then, the inclusion level was calculated as the abundance of the retained isoform divided by the abundance of both isoforms. The statistical method and codes from rMATS ^40^ were utilized to look for introns with significant inclusion level differences between wild-type and knockout cells. For each pair of compared groups, introns with a sum of the number of reads supporting spliced isoforms or retained isoforms less than 20 in either sample were filtered before statistical testing to remove potential false positives. The *P* values on the cumulative plots of inclusion level differences were calculated using two-sided Mann-Whitney tests.

### Statistics and reproducibility

Data are presented as the mean ± s.e.m. We used the paired Student’s t test for comparisons between the two experimental groups. All statistics were performed using GraphPad Prism 6 or R (4.2.1). The number of biological replicates for each experiment is indicated in the figure legends. At least four independent experiments of (e)RIP-PEN-seq were performed for both HEK293T-FLAG-15.5K and Hepa1-6-FLAG-15.5K cells. Two independent sets of HEK293T, HCT116, U-87 MG, C2C12 and Hepa1-6 RNA samples were used for CD-seq or sRNA-seq analysis. Two independent sets of human and mouse tissue samples were used for klRNA expression analysis, and three independent sets of HEK293T, HCT116, U-87 MG, C2C12 and Hepa1-6 RNA samples were used for klRNA expression validation. All agarose gels are representative of at least two biological replicates. No statistical method was used to predetermine the sample size.

## Data and code availability

The custom Perl and R scripts used in this study are available on request to the corresponding authors. All sequencing data that support the findings of this study have been deposited in NCBI’s Gene Expression Omnibus (GEO).

1. CD-seq in HEK293T, HCT116, U-87 MG, Hepa1-6 and C2C12 cells accession number: GSE134411 (https://www.ncbi.nlm.nih.gov/geo/query/acc.cgi?token=edmfyyiehvgdpgt&acc=GSE134411).
2. RIP-PEN-seq in HEK293T cells accession number: GSE160970 (https://www.ncbi.nlm.nih.gov/geo/query/acc.cgi?token=gzefamsgzjuthgp&acc=GSE160970).
3. RIP-PEN-seq in Hepa1-6 cells accession number: GSE182757 (https://www.ncbi.nlm.nih.gov/geo/query/acc.cgi?token=uvkxoguehhgzpex&acc=GSE182757).
4. PEN-seq in HCT116, U-87 MG, Hela, HEK293T, HepG2 and K562 accession number: GSE160887 (https://www.ncbi.nlm.nih.gov/geo/query/acc.cgi?token=mfobckqwflathed&acc=GSE160887).
5. PEN-seq in 15.5K knockdown HEK293T cells accession number: GSE186849 (https://www.ncbi.nlm.nih.gov/geo/query/acc.cgi?token=wtunaksafjkrxsv&acc=GSE186849).
6. PEN-seq in HEK293T and HCT116 cell fractions accession number: GSE182843 (https://www.ncbi.nlm.nih.gov/geo/query/acc.cgi?token=mtqzkamgzvmvdsn&acc=GSE182843).
7. RNA-seq in 15.5K knockdown HEK293T cells accession number: GSE182759 (https://www.ncbi.nlm.nih.gov/geo/query/acc.cgi?token=yvipyssapxannqz&acc=GSE182759).
8. SHAPE-MaP in HEK293T cells accession number: GSE220470 (https://www.ncbi.nlm.nih.gov/geo/query/acc.cgi?token=cpizwqeqzfupfef&acc=GSE220470

## Acknowledgments

We thank Dr. Lin Huang from Sun Yat-sen Memorial Hospital for his valuable suggestions on this manuscript. This work was supported by the National Key R&D Program of China (2024YFC3405901 (J.Y.), 2022YFA1303300 (J.Y.), 2024YFC3407001 (B.L.)); the National Natural Science Foundation of China (32225011 (J.Y.), 32430019 (J.Y.), 32370588 (B.L.), 32470598 (S.R.L.)); Special Project for Research and Development in Key areas of Guangdong Province (2024B1111130003) (J.Y.), Guangdong Basic and Applied Basic Research Foundation (2025B1515020051 (B.L.), 2025A1515010287 (S.R.L.)), Funding by Science and Technology Projects in Guangzhou (2025A04J3301 (B.L.), 2025A04J3498 (S.R.L.)).

## Author contributions

Conceptualization: JHY, BL, LHQ; Methodology: BL, QL, ARL, JHY; Investigation: BL, QL, ARL, HFG, SRL, WJZ, YLL, GW, JHH, KRZ, JHY; Visualization: BL, ARL, JHY; Funding acquisition: JHY, BL, SRL; Project administration: JHY, LHQ, BL; Supervision: JHY, LHQ, BL; Writing – original draft: BL, JHY, QL, ARL; Writing – review & editing: BL, JHY, LHQ.

## Extended Data Figure Legends

**Extended Data Fig. 1.**
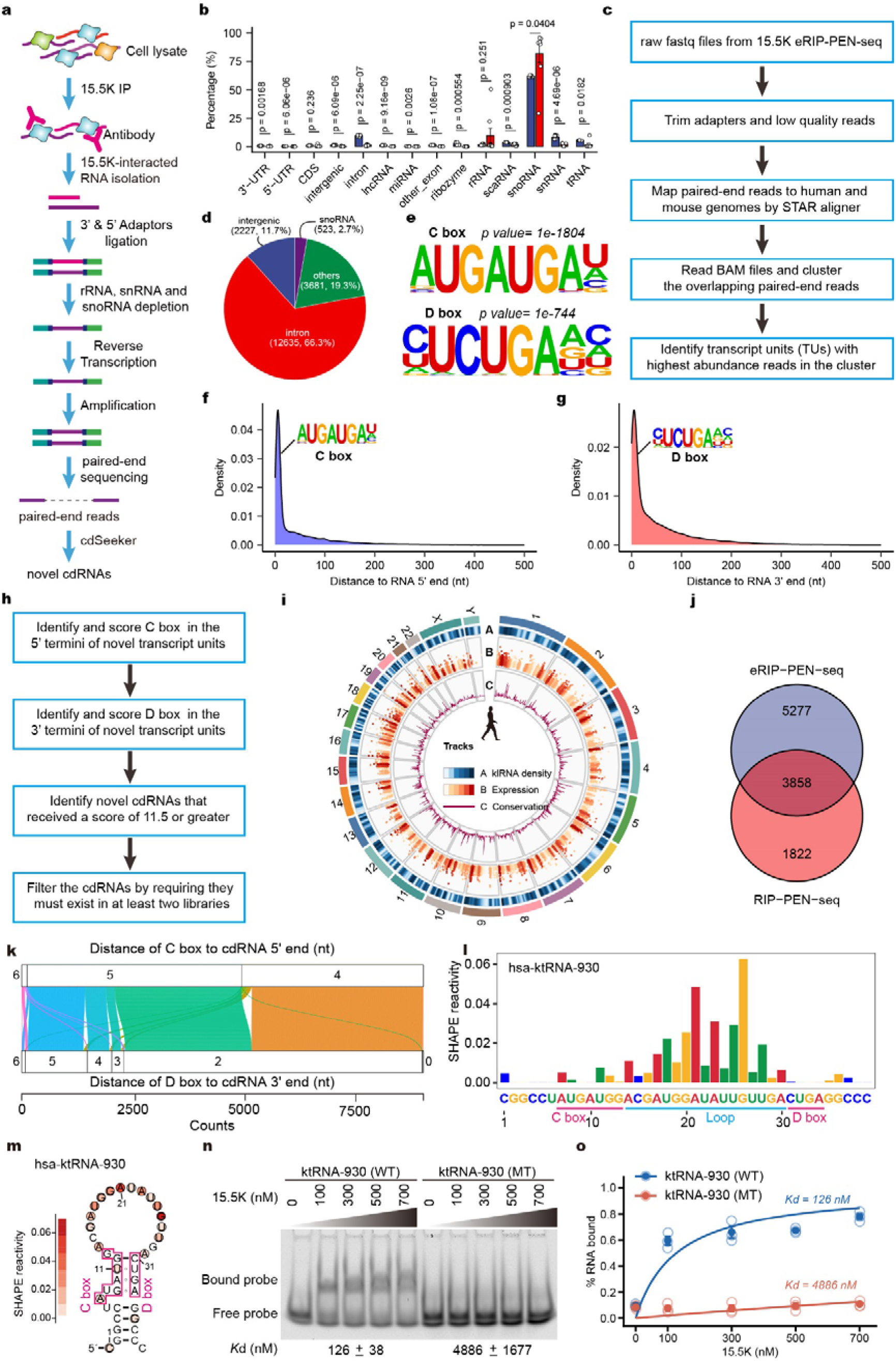
Identification and structural characterization of 15.5K-associated klRNAs in human cells. **a,** Schematic overview of enhanced RIP-PEN-seq (eRIP-PEN-seq) for profiling full-length RNAs associated with 15.5K. **b,** Comparison of RNA biotype composition between eRIP-PEN-seq and conventional RIP-PEN-seq libraries. Data are mean ± s.e.m (n = 4, biological replicates), two-tailed and paired t-test. **c,** Bioinformatic workflow for identification of novel transcript units (TUs). **d,** Genomic annotation of novel transcript units identified from 15.5K eRIP-PEN-seq. **e,** Enriched sequence motifs identified from novel transcript units, revealing canonical C-box and D-box motifs. **f, g,** Distribution of C-box (**f**) and D-box (**g**) positions relative to RNA termini. **h,** Computational workflow for identification of candidate cdRNAs. **i,** Genome-wide distribution, expression and evolutionary conservation of human cdRNAs identified by eRIP-PEN-seq. RPM: reads per million reads. The plot legend is shown in the right panel. **j,** Comparison of cdRNAs identified from eRIP-PEN-seq and RIP-PEN-seq in HEK293T cells. **k,** Alluvial plot of the distance of C box and D box to cdRNA ends. **l,** The SHAPE reactivity signal on hsa-ktRNA-930. The C and D boxes and the loop region are marked with pink and blue underlines in the bar plot, respectively. The SHAPE reactivity is calculated from merged n = 4 biological replicates. **m,** The predicted secondary structure of the K-turn on hsa-ktRNA-930. The C and D boxes are indicated with pink boxes in the structure figures. The SHAPE reactivity is calculated from merged n = 4 biological replicates. **n, o,** RNA electrophoretic mobility shift assay (REMSA) (**n**) was used to assess the binding of recombinant 15.5K and the wild-type (WT) or D box-mutant (MT) ktRNA-930 RNA probes. Quantification of independent REMSA (**o**) reveals the percentage of bound probes with increasing concentration of 15.5K over three replicates for WT- and MT-ktRNA-930, respectively. The binding curves are presented as mean ± s.d. (n = 3 for each data point). The dissociation constant (Kd, nM) values were indicated at the lower panel of the REMSA image.

**Extended Data Fig. 2.**
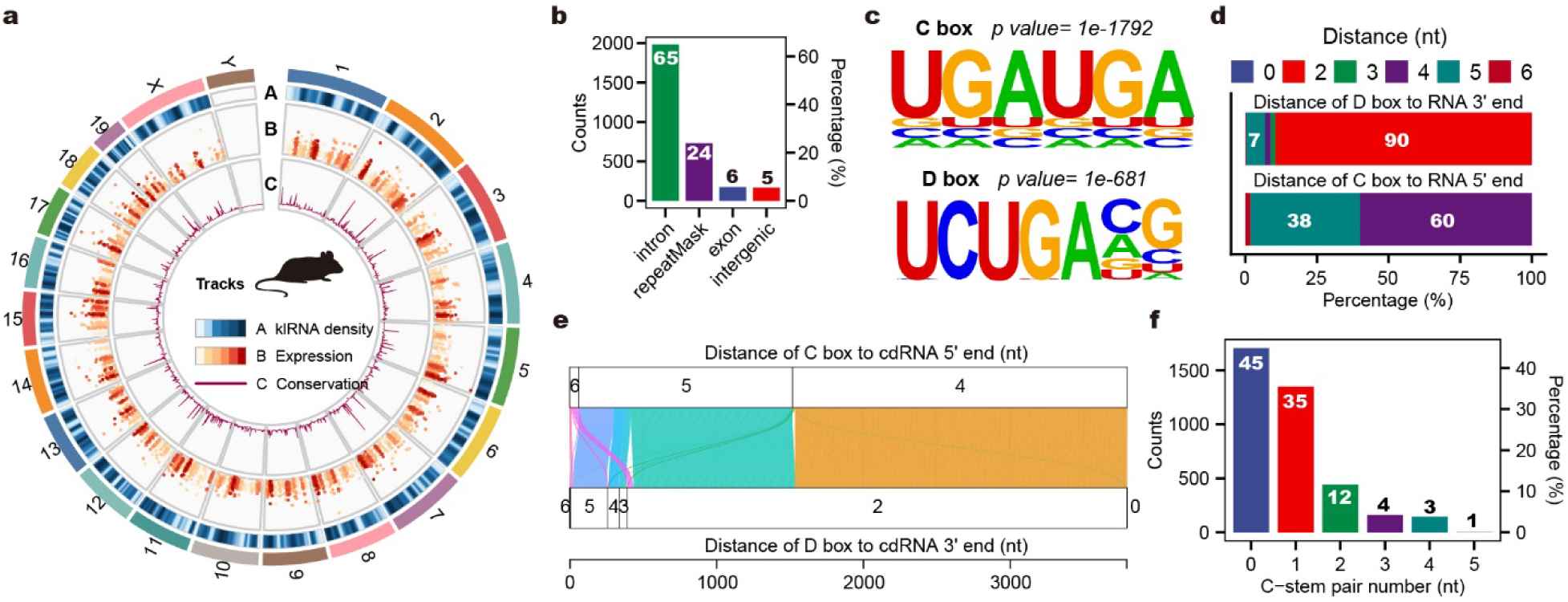
Genome-wide identification of klRNAs in mouse cells by 15.5K RIP-PEN-seq. **a,** Genome-wide distribution, expression and evolutionary conservation of mouse cdRNAs. **b,** Distribution and proportion of mouse cdRNAs across genomic annotation categories. **c,** Enriched C-box and D-box motifs identified from mouse cdRNAs. **d,** Percentage of cdRNAs at different distances from the RNA end for C box and D box. **e,** Alluvial plot of the distance of C box and D box to cdRNA ends. **f,** Statistics on the number and proportion of base pairs in the C-stem.

**Extended Data Fig. 3.**
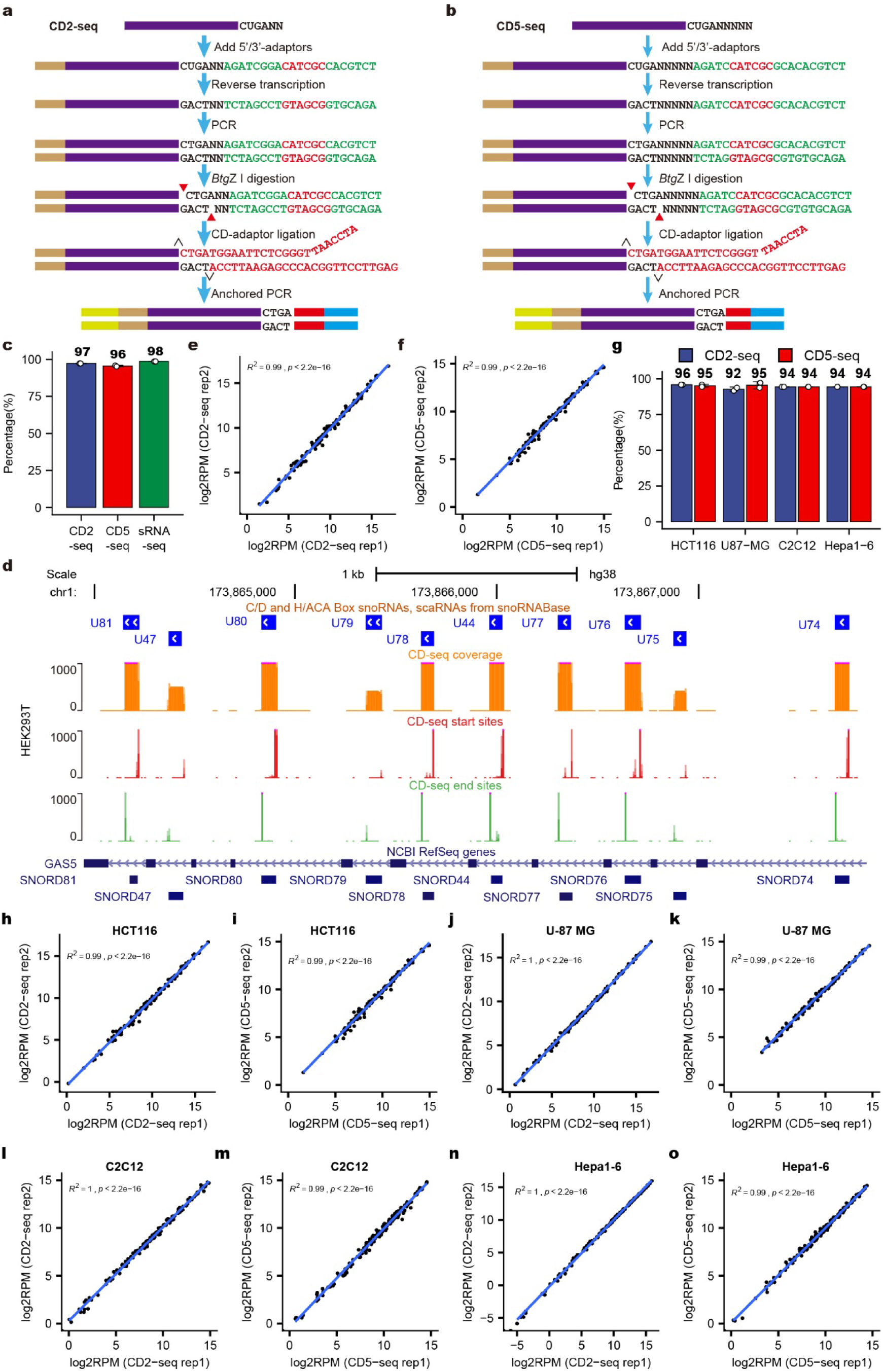
Performance evaluation and validation of CD-seq for full-length box C/D RNA profiling. **a, b,** Detailed workflows of CD2-seq (**a**) and CD5-seq (**b**). **c,** Detection sensitivity of canonical box C/D RNAs by CD-seq compared with sRNA-seq. **d,** CD2-seq and CD5-seq profiles of canonical box C/D snoRNAs (coverage, orange; 5’-start, red; 3’-end, green) located within the introns of the GAS5 gene. **e, f,** Reproducibility of CD2-seq (**e**) and CD5-seq (**f**) measurements between biological replicates. The correlation of the reads was calculated with the Pearson correlation coefficient. **g,** Detection of canonical box C/D RNAs across multiple mammalian cell types. **h, i,** Scatter plots of canonical box C/D RNA expression computed from CD2-seq (**h**) and CD5-seq (**i**) libraries generated from biological replicates of HCT116 cells. The correlation of the reads was calculated with the Pearson correlation coefficient. **j, k,** Scatter plots of canonical box C/D RNA expression computed from the CD2-seq (**j**) and CD5-seq (**k**) libraries generated from biological replicates of U-87 MG cells. The correlation of the reads was calculated with the Pearson correlation coefficient. **l, m,** Scatter plots of canonical box C/D RNA expression computed from the CD2-seq (**l**) and CD5-seq (**m**) libraries generated from biological replicates of C2C12 cells. The correlation of the reads was calculated with the Pearson correlation coefficient. **n, o,** Scatter plots of canonical box C/D RNA expression computed from the CD2-seq (**n**) and CD5-seq (**o**) libraries generated from biological replicates of Hepa1-6 cells. The correlation of the reads was calculated with the Pearson correlation coefficient.

**Extended Data Fig. 4.**
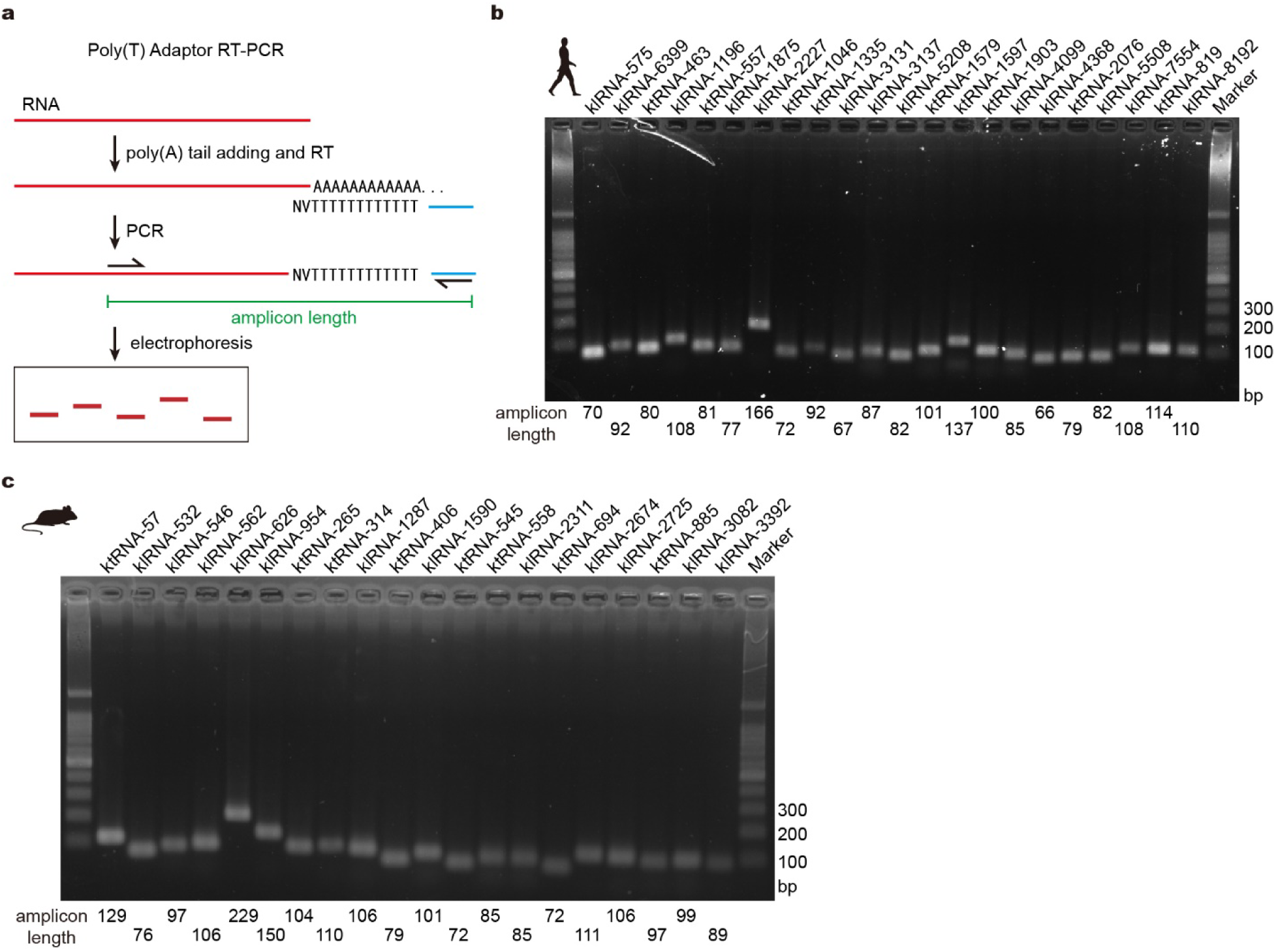
Experimental validation of human and mouse klRNAs by poly(T)-adaptor RT-PCR. **a,** Workflow for determining RNA 3’ termini using poly(T)-adaptor RT-PCR. **b,** Validation of 22 selected human klRNAs and ktRNAs through poly(T) adaptor RT-PCR analysis of total RNA extracted from human HEK293T cells. The length of the amplicons is indicated below each lane. **c,** Validation of 20 selected mouse klRNAs and ktRNAs using poly(T) adaptor RT-PCR in total RNA extracted from mouse C2C12 cells. The lengths of the amplicons are indicated below each lane.

**Extended Data Fig. 5.**
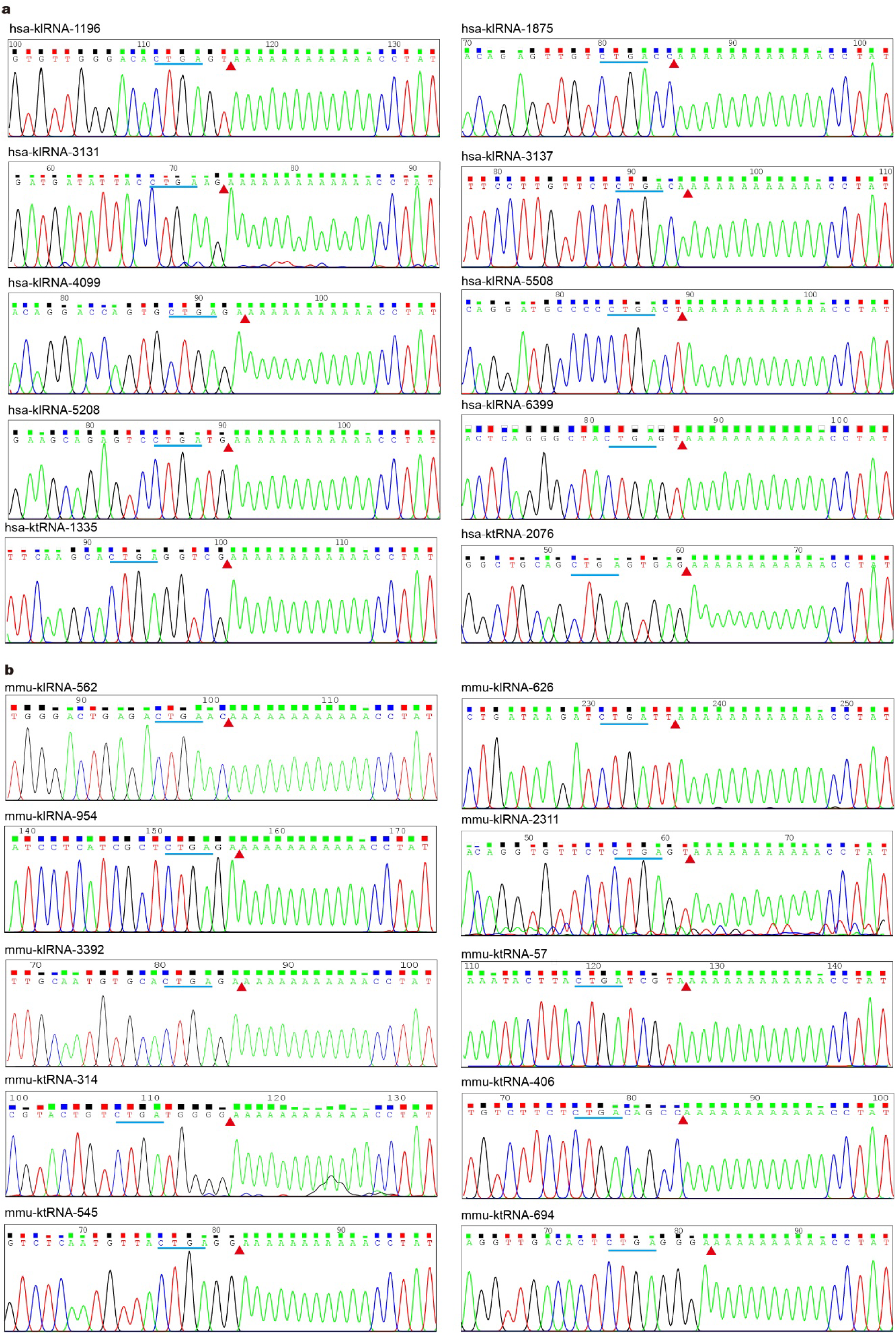
Sanger validation of precise 3’ termini of klRNAs and ktRNAs. **a, b,** Sanger sequencing confirms the precise 3’ termini of representative human and mouse klRNAs and ktRNAs. The conserved D-box motif is indicated, and the experimentally determined RNA termini are marked with a light red triangle.

**Extended Data Fig. 6.**
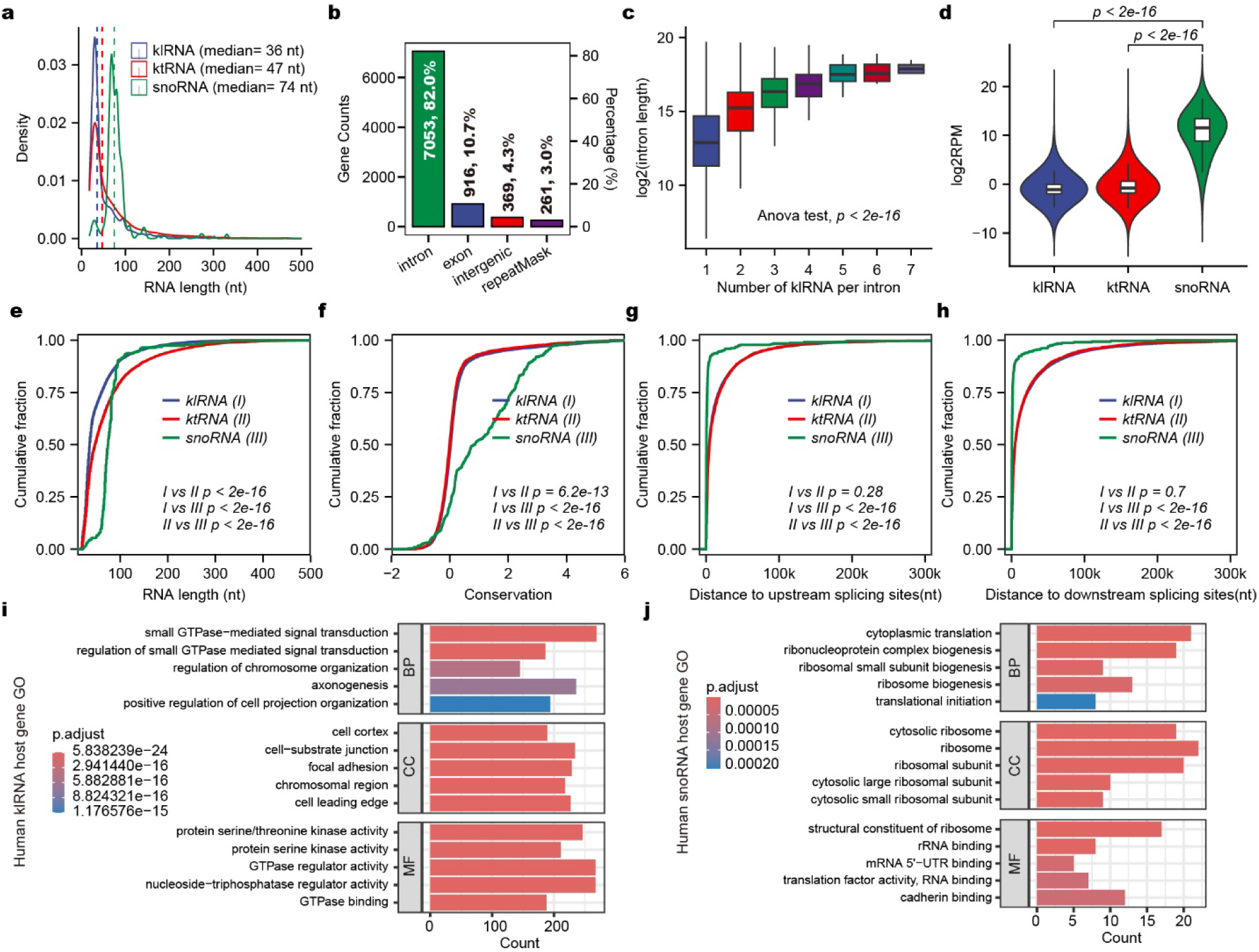
Genomic, structural and expression characteristics of human klRNAs. **a,** Length distribution of canonical box C/D snoRNAs, ktRNAs and klRNAs. The dashed lines indicate the median values for the different groups. **b,** Counts and percentages of klRNA□associated genes across distinct genomic annotation categories. **c,** Relationship between intron length and the number of encoded klRNAs. Box plots show minimum value, first quartile, median, third quartile and maximum value. The one-way ANOVA test was used to calculate the P values. **d,** Expression distribution of different classes of structured RNAs. Box plots show minimum value, first quartile, median, third quartile and maximum value. RPM: reads per million reads. The Mann-Whitney-Wilcoxon test was used to calculate the P values of the differences between the two categories. **e,** Cumulative fraction (CDF) plot showing RNA length between canonical box C/D snoRNAs, ktRNAs, and klRNAs identified in humans. The Mann-Whitney-Wilcoxon test was used to calculate the P values of the differences in length between the two categories. **f,** Cumulative fraction (CDF) plot showing RNA sequence conservation between canonical box C/D RNAs, ktRNAs and klRNAs identified in humans. The Mann-Whitney-Wilcoxon test was used to calculate the P values of the differences in conservation between the two categories. **g, h,** Cumulative curves of both upstream (**g**) and downstream (**h**) distances for canonical box C/D snoRNAs, ktRNAs, and klRNAs in humans. The Mann-Whitney-Wilcoxon test was used to calculate the P values of the differences in distance between the two categories. **i, j,** Gene Ontology (GO) enrichment analysis of klRNAs (**i**) and the canonical box C/D snoRNA (**j**) host genes within humans. The top five enriched GO categories (BP, CC and MF) are listed. The P values (p.adjust) were sorted by significance (low, blue; high, red). BP, Biological Process; CC, Cellular Compartment; MF, Molecular Function.

**Extended Data Fig. 7.**
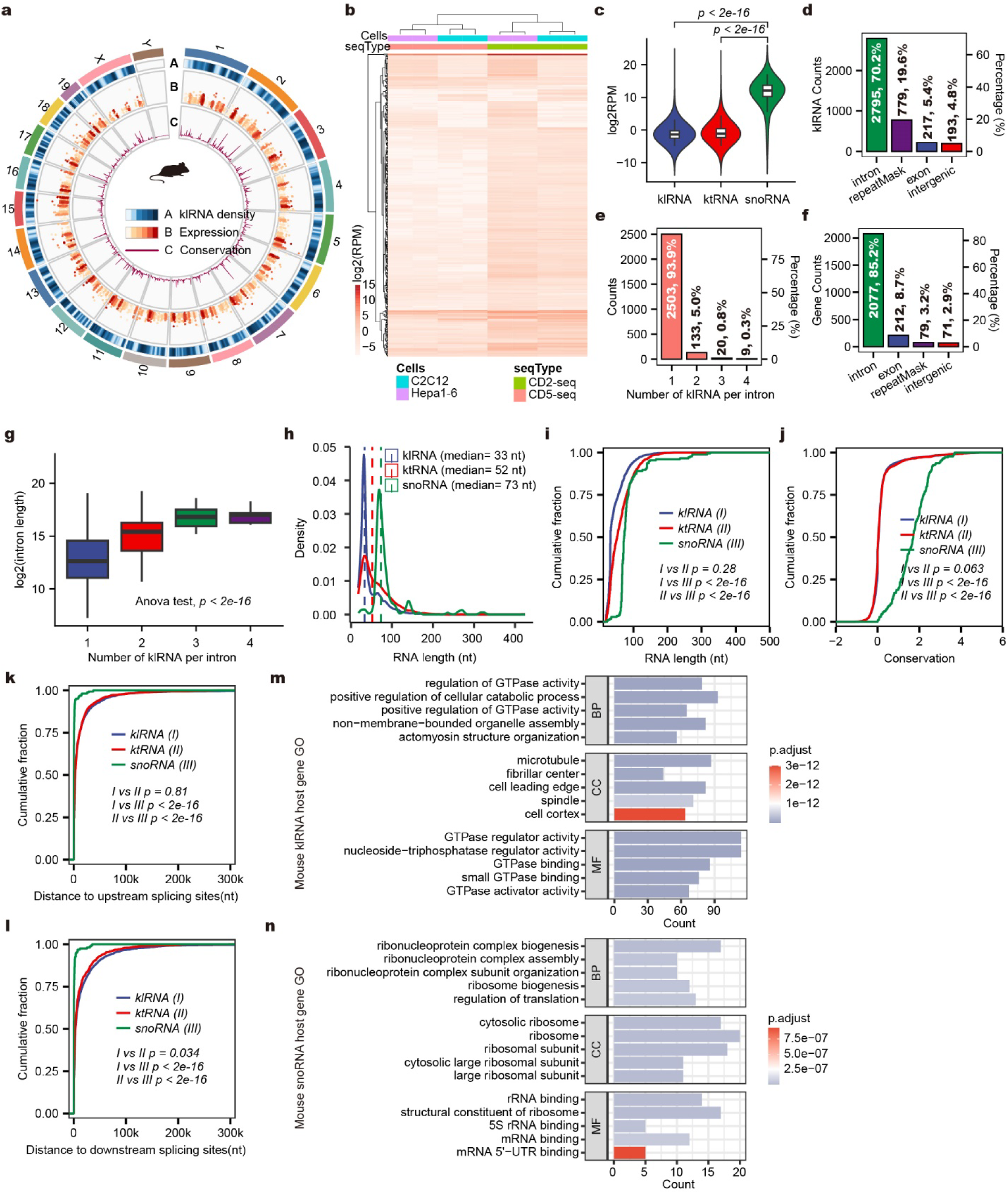
Genomic, structural and expression characteristics of mouse klRNAs. **a,** Genome-wide distribution, expression and conservation of mouse klRNAs. RPM: reads per million reads. The plot legend is shown in the lower panel. **b,** Heatmap of mouse klRNAs expression in C2C12 and Hepa1-6 CD-seq libraries. **c,** Expression comparison among box C/D snoRNAs, ktRNAs and klRNAs. Box plots show minimum value, first quartile, median, third quartile and maximum value. RPM: reads per million reads. The Mann-Whitney-Wilcoxon test was used to calculate the P values of the differences in expression level between the two categories. **d,** Numbers and percentage of mouse klRNAs in different genomic annotation categories. **e,** The count of introns encoding varying quantities of klRNAs. **f,** Counts and percentages of klRNA associated genes across distinct genomic annotation categories. **g,** Box plots of distributions of length for introns encoding varying quantities of klRNAs. Box plots show minimum value, first quartile, median, third quartile and maximum value. The one-way ANOVA test was used to calculate the P values. **h,** Density plot comparing the length distribution between the canonical box C/D snoRNAs, ktRNAs and klRNAs in mice. The dashed lines indicate the median values for the different groups. **i,** Cumulative fraction (CDF) plot showing RNA length between canonical box C/D snoRNAs, ktRNAs, and klRNAs identified in mice. The Mann-Whitney-Wilcoxon test was used to calculate the P values of the differences in length between the two categories. **j,** Cumulative fraction (CDF) plot showing RNA sequence conservation between canonical box C/D snoRNAs, ktRNAs, and klRNAs identified in mice. The Mann-Whitney-Wilcoxon test was used to calculate the P values of the differences in conservation between the two categories. **k, l,** Cumulative curves of both upstream (**k**) and downstream (**l**) distances for canonical box C/D snoRNAs, ktRNAs, and klRNAs in mice. The Mann-Whitney-Wilcoxon test was used to calculate the P values of the differences in distance between the two categories. **m, n,** Gene Ontology (GO) enrichment analysis of klRNAs (**m**) and the canonical box C/D snoRNA (**n**) host genes within mice. The top five enriched GO categories (BP, CC and MF) are listed. The P values (p.adjust) were sorted by significance (low, blue; high, red). BP, Biological Process; CC, Cellular Compartment; MF, Molecular Function.

**Extended Data Fig. 8.**
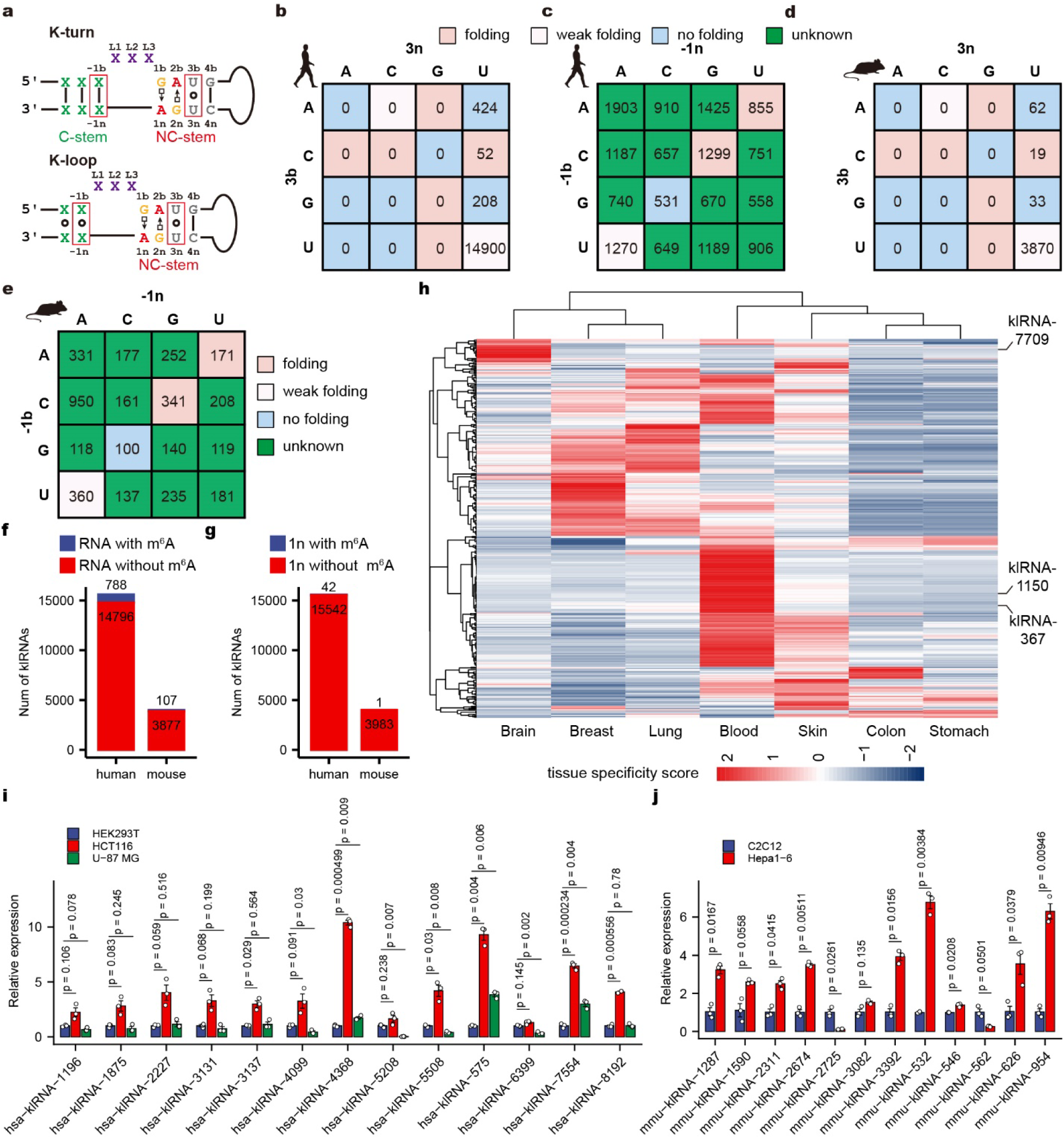
Sequence features, structural constraints and expression patterns of klRNAs. **a,** Consensus secondary structures of canonical K-turns and K-loops. The nucleotide positions in the K-turn structure are named according to the nomenclature rules. **b,** Matrix plot showing the number of human klRNAs with the indicated nucleotide in the 3b:3n sequences. **c,** Number of human klRNAs with the four possible Watson-Crick base pairs in the -1b:-1n position. **d,** Matrix plot showing the number of mouse klRNAs with the indicated nucleotide in the 3b:3n sequences. **e,** Number of mouse klRNAs with the four possible Watson-Crick base pairs in the -1b:-1n position. **f,** Number of klRNAs with or without m^6^A modification in humans and mice. **g,** Number of klRNAs with or without m^6^A modification at the 1n position in humans and mice. **h,** Tissue-specific expression profiles of human klRNAs. The expression levels of klRNAs are displayed in the rows and the tissues are shown in the columns. The rows and columns are sorted based on k-means clustering of klRNAs genes. The color intensity represents the tissue-specific score (JS score) as calculated for each klRNA using the csSpecificity function in the CummeRbund R package. Representative klRNAs are indicated in the right panel. **i,** Relative expression analysis of klRNAs in HEK293T, HCT116 and U-87 MG cells through poly(T) adaptor-based qPCR. Data are presented as the fold change (mean ± s.e.m, n=3) normalized to HEK293T. U6 RNA served as a reference gene. The P values were calculated using a two-sided t test. **j,** Relative expression analysis of klRNAs in C2C12 and Hepa1-6 cells through poly(T) adaptor-based qPCR. Data are presented as the fold change (mean ± s.e.m, n=3) normalized to C2C12. U6 RNA served as a reference gene. The P values were calculated using a two-sided t test.

**Extended Data Fig. 9.**
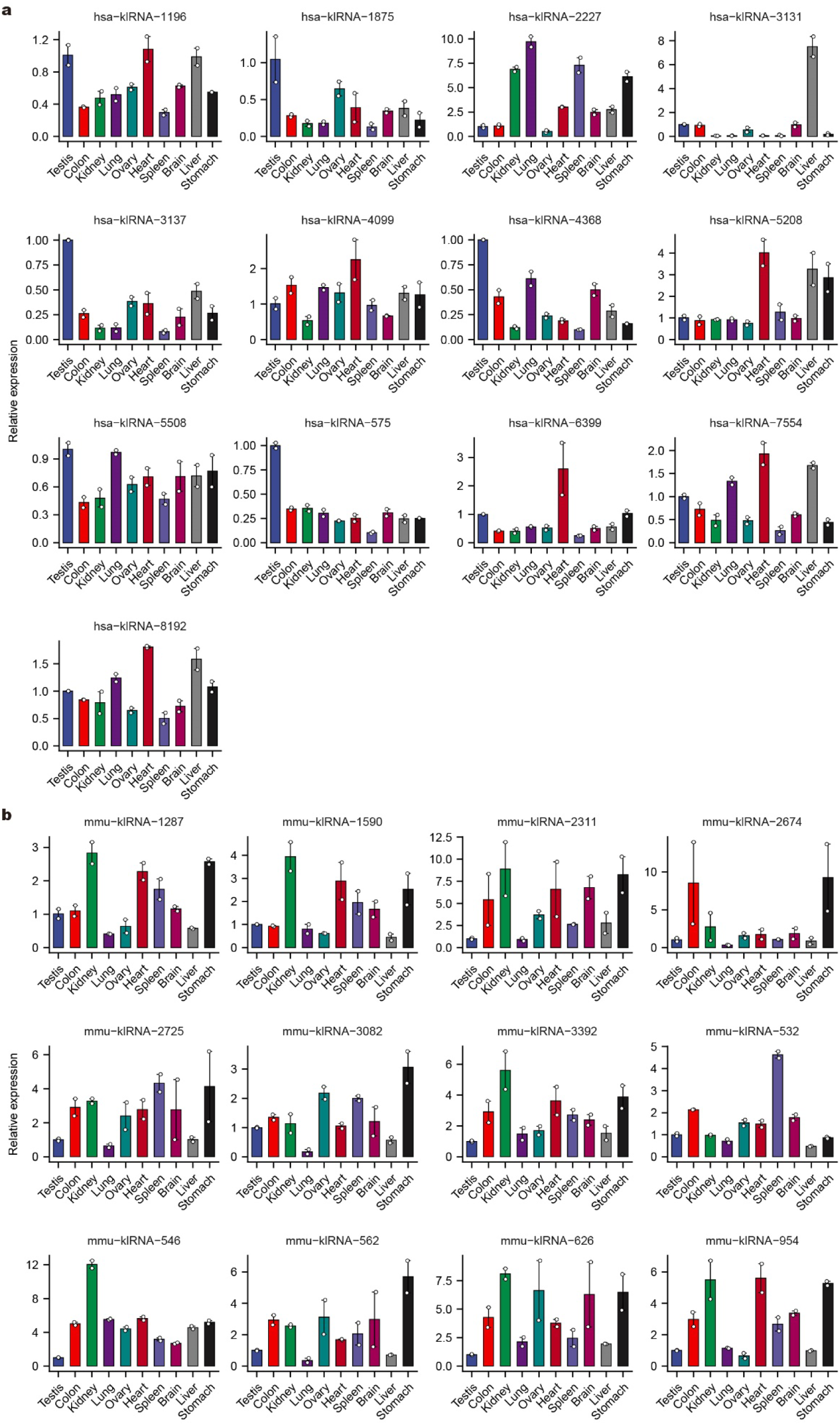
Tissue expression profiles of human and mouse klRNAs. **a, b,** Relative expression analysis of representative klRNAs in human (**a**) and mouse (**b**) tissues through poly(T) adaptor-based qPCR. Data are presented as the fold change (mean ± s.e.m, n=2) normalized to the testis. U6 RNA served as a reference gene.

**Extended Data Fig. 10.**
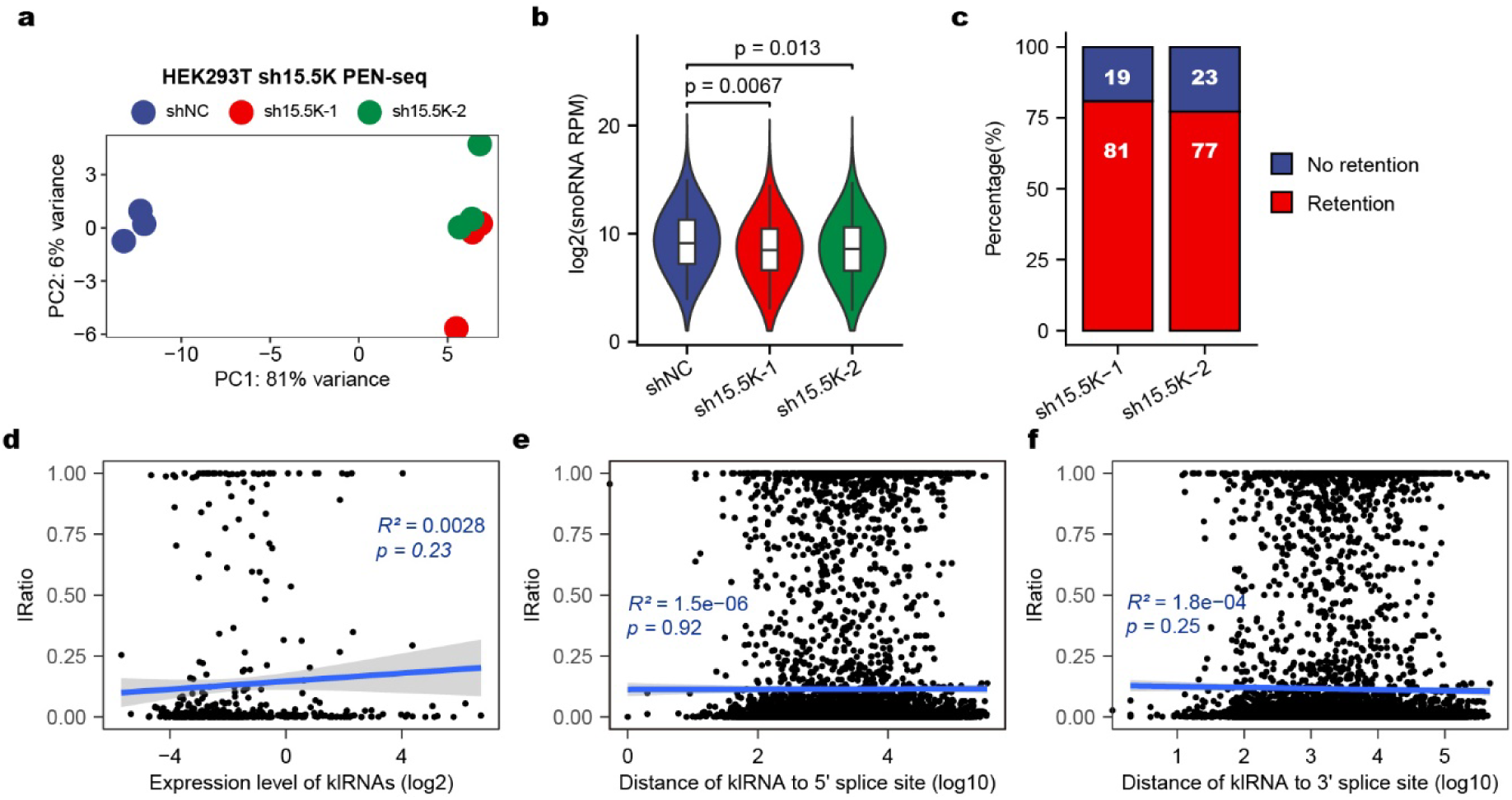
Transcriptome-wide effects of 15.5K depletion on klRNA expression and intron retention. **a,** PCA was performed based on PEN-seq profiles to illustrate the overall similarity and clustering of HEK293T-sh15.5K cell samples. **b,** Global expression distribution of canonical box C/D snoRNAs following 15.5K depletion. Box plots show minimum value, first quartile, median, third quartile and maximum value. RPM: reads per million reads. The P-values of the differences in distance between the two categories were determined by the Mann-Whitney-Wilcoxon test. **c,** Fraction of significantly retained introns (filtered by P <0.05) identified after 15.5K knockdown. **d,** Correlation analysis between klRNA expression levels and the retention rates of their host introns. **e,** Correlation analysis between klRNA distance to intronic 5’ splice sites and the retention rates of their host introns. **f,** Correlation analysis between klRNA distance to intronic 3’ splice sites and the retention rates of their host introns.

**Extended Data Fig. 11.**
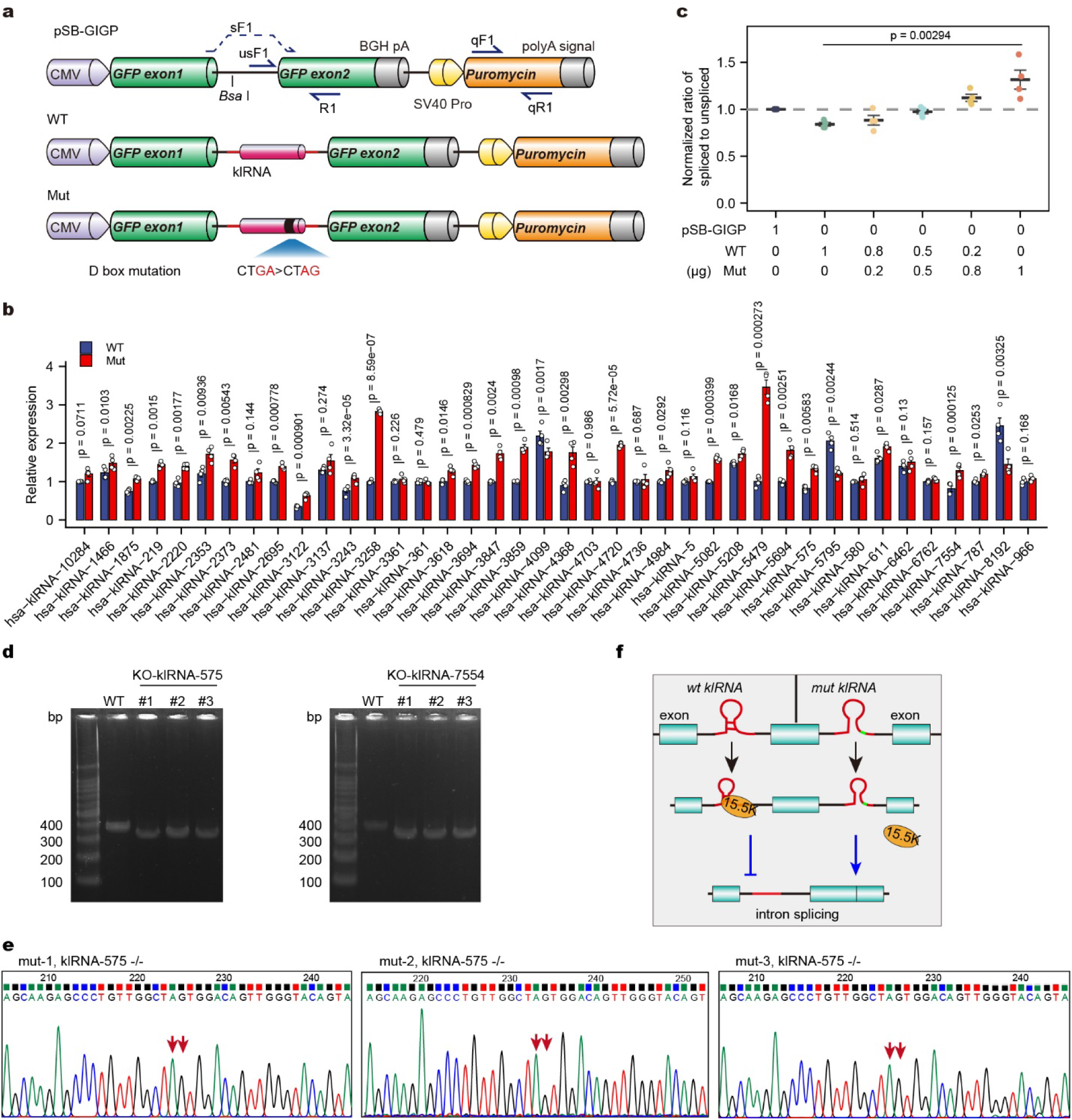
Functional validation of klRNA-mediated autoregulation of host intron splicing. **a,** Schematic of the GFP splicing reporter design, klRNA cloning, and primers used for splicing efficiency detection. We used the primer pairs sF1/R1 and usF1/R1 for detecting mature and precursor GFP, respectively. The analysis of Puromycin with primer pairs qF1/qR1 served as internal control. **b,** qPCR analysis of the ratio of spliced to unspliced GFP RNA in HEK293T cells transfected with wild-type and mutant klRNAs (mutated from CTGA to CTAG). Data are mean ± s.e.m (n = 4, biological replicates), two-tailed and paired t-test. **c,** Dose-dependent effect of wild-type and mutant klRNA reporters on splicing regulation. Data are mean ± s.e.m (n = 4, biological replicates). The P-values among the indicated categories were determined by the one-way ANOVA. **d,** PCR analyses of genomic DNA obtained from klRNA-575 and klRNA-7554 knockout cells to validate biallelic klRNA deletion clones. **e,** Sanger DNA sequencing analysis of klRNA-575 genotype in Prime editing cells. The red arrows represent the edited bases. **f,** Proposed model for klRNA-mediated autoregulation of host intron splicing. Terminal C/D motif-derived K-loop structures recruit 15.5K to locally suppress excision of the host intron, whereas disruption of the K-loop abolishes 15.5K recruitment and relieves local splicing repression.

## References

1. Venter, J.C. et al. The sequence of the human genome. Science (New York, N.Y.) 291, 1304–1351 (2001).

2. Lander, E.S. et al. Initial sequencing and analysis of the human genome. Nature 409, 860–921 (2001).

3. Chen, L.L. & Kim, V.N. Small and long non-coding RNAs: Past, present, and future. Cell 187, 6451–6485 (2024).

4. Goldrich, M.J. et al. Widespread DNA off-targeting confounds RNA chromatin occupancy studies. Nat Biotechnol (2026).

5. Rearick, D. et al. Critical association of ncRNA with introns. Nucleic Acids Res. 39, 2357–2366 (2010).

6. Klein, D.J., Schmeing, T.M., Moore, P.B. & Steitz, T.A. The kink-turn: a new RNA secondary structure motif. EMBO J 20, 4214–4221 (2001).

7. Lilley, D.M. The K-turn motif in riboswitches and other RNA species. Biochim Biophys Acta 1839, 995–1004 (2014).

8. Schroeder, K.T., McPhee, S.A., Ouellet, J. & Lilley, D.M. A structural database for k-turn motifs in RNA. RNA 16, 1463–1468 (2010).

9. Nolivos, S., Carpousis, A.J. & Clouet-d’Orval, B. The K-loop, a general feature of the Pyrococcus C/D guide RNAs, is an RNA structural motif related to the K-turn. Nucleic Acids Res. 33, 6507–6514 (2005).

10. Li, B. et al. RIP-PEN-seq identifies a class of kink-turn RNAs as splicing regulators. Nature Biotechnology 42, 119–131 (2024).

11. Yang, J.H. et al. snoSeeker: an advanced computational package for screening of guide and orphan snoRNA genes in the human genome. Nucleic Acids Res 34, 5112–5123 (2006).

12. Henras, A.K., Dez, C. & Henry, Y. RNA structure and function in C/D and H/ACA s(no)RNPs. Curr Opin Struct Biol 14, 335–343 (2004).

13. Shi, R., Sun, Y.H., Zhang, X.H. & Chiang, V.L. Poly(T) adaptor RT-PCR. Methods Mol Biol 822, 53–66 (2012).

14. Consortium, E.P. An integrated encyclopedia of DNA elements in the human genome. Nature 489, 57–74 (2012).

15. Huang, L. et al. Structure and folding of four putative kink turns identified in structured RNA species in a test of structural prediction rules. Nucleic Acids Res 49, 5916–5924 (2021).

16. Huang, L., Wang, J. & Lilley, D.M. A critical base pair in k-turns determines the conformational class adopted, and correlates with biological function. Nucleic Acids Res 44, 5390–5398 (2016).

17. McPhee, S.A., Huang, L. & Lilley, D.M. A critical base pair in k-turns that confers folding characteristics and correlates with biological function. Nat Commun 5, 5127 (2014).

18. Liu, J. & Lilley, D.M. The role of specific 2’-hydroxyl groups in the stabilization of the folded conformation of kink-turn RNA. RNA 13, 200–210 (2007).

19. Lestrade, L. & Weber, M.J. snoRNA-LBME-db, a comprehensive database of human H/ACA and C/D box snoRNAs. Nucleic Acids Res 34, D158–162 (2006).

20. Bradley, R.K. & Anczukow, O. RNA splicing dysregulation and the hallmarks of cancer. Nat Rev Cancer 23, 135–155 (2023).

21. Zhang, Y. et al. Overexpression of lncRNAs with endogenous lengths and functions using a lncRNA delivery system based on transposon. J Nanobiotechnology 19, 303 (2021).

22. Chomczynski, P. & Sacchi, N. The single-step method of RNA isolation by acid guanidinium thiocyanate-phenol-chloroform extraction: twenty-something years on. Nat Protoc 1, 581–585 (2006).

23. Giraldez, M.D. et al. Comprehensive multi-center assessment of small RNA-seq methods for quantitative miRNA profiling. Nat Biotechnol 36, 746–757 (2018).

24. Han, E.S. et al. RecJ exonuclease: substrates, products and interaction with SSB. Nucleic Acids Res 34, 1084–1091 (2006).

25. Martin, M. Cutadapt removes adapter sequences from high-throughput sequencing reads. EMBnet.journal 17, 10–12 (2011).

26. Dobin, A. et al. STAR: ultrafast universal RNA-seq aligner. Bioinformatics 29, 15–21 (2013).

27. Haeussler, M. et al. The UCSC Genome Browser database: 2019 update. Nucleic Acids Res 47, D853–D858 (2019).

28. Frankish, A. et al. GENCODE reference annotation for the human and mouse genomes. Nucleic Acids Res 47, D766–D773 (2019).

29. Xie, F. et al. deepBase v3.0: expression atlas and interactive analysis of ncRNAs from thousands of deep-sequencing data. Nucleic Acids Res 49, D877–D883 (2021).

30. Jorjani, H. et al. An updated human snoRNAome. Nucleic Acids Res 44, 5068–5082 (2016).

31. Kishore, S. et al. Insights into snoRNA biogenesis and processing from PAR-CLIP of snoRNA core proteins and small RNA sequencing. Genome Biol 14, R45 (2013).

32. Pruitt, K.D. et al. RefSeq: an update on mammalian reference sequences. Nucleic Acids Res 42, D756–763 (2014).

33. Quinlan, A.R. & Hall, I.M. BEDTools: a flexible suite of utilities for comparing genomic features. Bioinformatics 26, 841–842 (2010).

34. Yu, G., Wang, L.G., Han, Y. & He, Q.Y. clusterProfiler: an R package for comparing biological themes among gene clusters. OMICS 16, 284–287 (2012).

35. Magoc, T. & Salzberg, S.L. FLASH: fast length adjustment of short reads to improve genome assemblies. Bioinformatics 27, 2957–2963 (2011).

36. Busan, S. & Weeks, K.M. Accurate detection of chemical modifications in RNA by mutational profiling (MaP) with ShapeMapper 2. RNA 24, 143–148 (2018).

37. Sieg, J.P., Forstmeier, P.C. & Bevilacqua, P.C. JPSieg/R2easyR: R2easyR v1.1 (Version v1.1). Zenodo. 10.5281/zenodo.4683742 (2021).

38. Weinberg, Z. & Breaker, R.R. R2R--software to speed the depiction of aesthetic consensus RNA secondary structures. BMC Bioinformatics 12, 3 (2011).

39. Love, M.I., Huber, W. & Anders, S. Moderated estimation of fold change and dispersion for RNA-seq data with DESeq2. Genome Biol 15, 550 (2014).

40. Shen, S. et al. rMATS: robust and flexible detection of differential alternative splicing from replicate RNA-Seq data. P Natl Acad Sci USA 111, E5593–E5601 (2014).

